# Influence of stereochemistry in a local approach for calculating protein conformations

**DOI:** 10.1101/2024.09.18.613236

**Authors:** Wagner Da Rocha, Leo Liberti, Antonio Mucherino, Thérèse E. Malliavin

## Abstract

Protein structure prediction is usually based on the use of local conformational information coupled with long-range distance restraints. Such restraints can be derived from the knowledge of a template structure or the analysis of protein sequence alignment in the framework of models arising from the physics of disordered systems. The accuracy of approaches based on sequence alignment, however, is limited in the case where the number of aligned sequences is small. Here we derive protein conformations using only local conformations knowledge by means of the interval Branch-and-Prune algorithm. The computation efficiency is directly related to the knowledge of stereochemistry (bond angle and *ω* values) along the protein sequence, and in particular to the variations of the torsion angle *ω*. The impact of stereochemistry variations is particularly strong in the case of protein topologies defined from numerous long-range restraints, as in the case of protein of *β* secondary structures. The systematic enumeration of the conformations improves the efficiency of the calculations. The analysis of DNA codons permits to connect the variations of torsion angle *ω* to the positions of rare DNA codons.

## Introduction

The approaches for predicting protein structures from the knowledge of their primary sequence have undergone enormous developments during the last decades. ^1–3^ One of the most recent steps of this progress is the use of deep learning approaches. ^4–7^ These *in silico* predictions pave the way towards protein function prediction and drug design and can be thus considered as founding steps towards a reasoned interference with physiological processes, health problems, or plant engineering.

In the domain of protein structure prediction, template-free *in silico* approaches uses local structural information coupled to long-range proximities.^8^ The relative importance of these two pieces of information is essential for a successful prediction, as pointed out by Skolnick et al^9^ already long ago. As the development of covariance approaches for multiple sequence alignments^10–12^ permits the prediction of long-range restraints, a consensus was found on the fact that prediction methods must be based on the local and long-range pieces of information.^13^

The recently flourishing deep learning approaches^4–7^ have followed the same path, capitalizing on the availability of huge databases of protein structures and sequences. ^14,15^ The success of all prediction methods is thus quite dependent on the availability of long-range restraints and consequently on the availability of multiple sequence alignment. Prediction methods for the torsion angles *ϕ* and *ψ*, however, may rely on a unique protein sequence.^16–20^ Consequently, local structure prediction can be inferred independently of alignment information.

In several cases, long-range proximity information cannot be obtained because the size of the corresponding sequence alignments is insufficient. An obvious case arises in the presence of disordered regions involving many conformations, which prevents the determination of precise proximities. Besides, for some protein families, the number of aligned sequences is too small for statistically determining the long-range restraints.^21^ Proteins for which expression frameshift conducts to the expression of various polypeptides are also cases where the multiple sequence alignment does not provide reliable information. ^22^

We investigate here whether local structure information is sufficient to determine the protein fold. Of course, local and global structural pieces of information are closely linked: we are aware of the artificial nature of their separation. The present work should be considered a geometric investigation of the relative importance of local and global information for calculating protein conformations. In a previous analysis,^23^ it was shown that the use of distances restraints based on local geometry permitted to calculate protein conformations closer to the target protein structures. In addition, some initial investigations in line with the present work have been conducted.^24^

For our purpose, we employ a purely geometric approach, the interval Branch-and-Prune (iBP) algorithm, proposed some years ago to solve the problem of distance geometry in the frame of protein structure.^25–28^ The adaptation of iBP to intrinsically disordered proteins and regions is known as Threading-Augmented interval Branch-and-Prune (TAiBP).^28,29^ It systematically enumerates protein conformations while heuristically overcoming the intrinsic combinatorial barrier. Since then, TAiBP has been shown to allow the analysis of the conformational space of various flexible or disordered proteins.^30–32^

In the present work, we test several variants of the iBP algorithm with different levels of knowledge of the local geometry information (Table S1), on a database of 308 protein structures smaller than 100 residues (Figure 1B). These high-resolution X-ray crystallographic structures were selected in particular because they contain at least two secondary structural elements, *α* helices or *β* strands. In the following, the torsion angles *ϕ* and *ψ* will be assumed to be known within 5*^◦^* intervals (or within 40*^◦^* for loops in enumerating iBP runs), and the focus will be put on the variations of the torsion angle *ω* and of the bond angles. Larger variations of *ϕ* and *ψ* can be in principle taken into account using TAiBP.^29^

**Figure 1:**
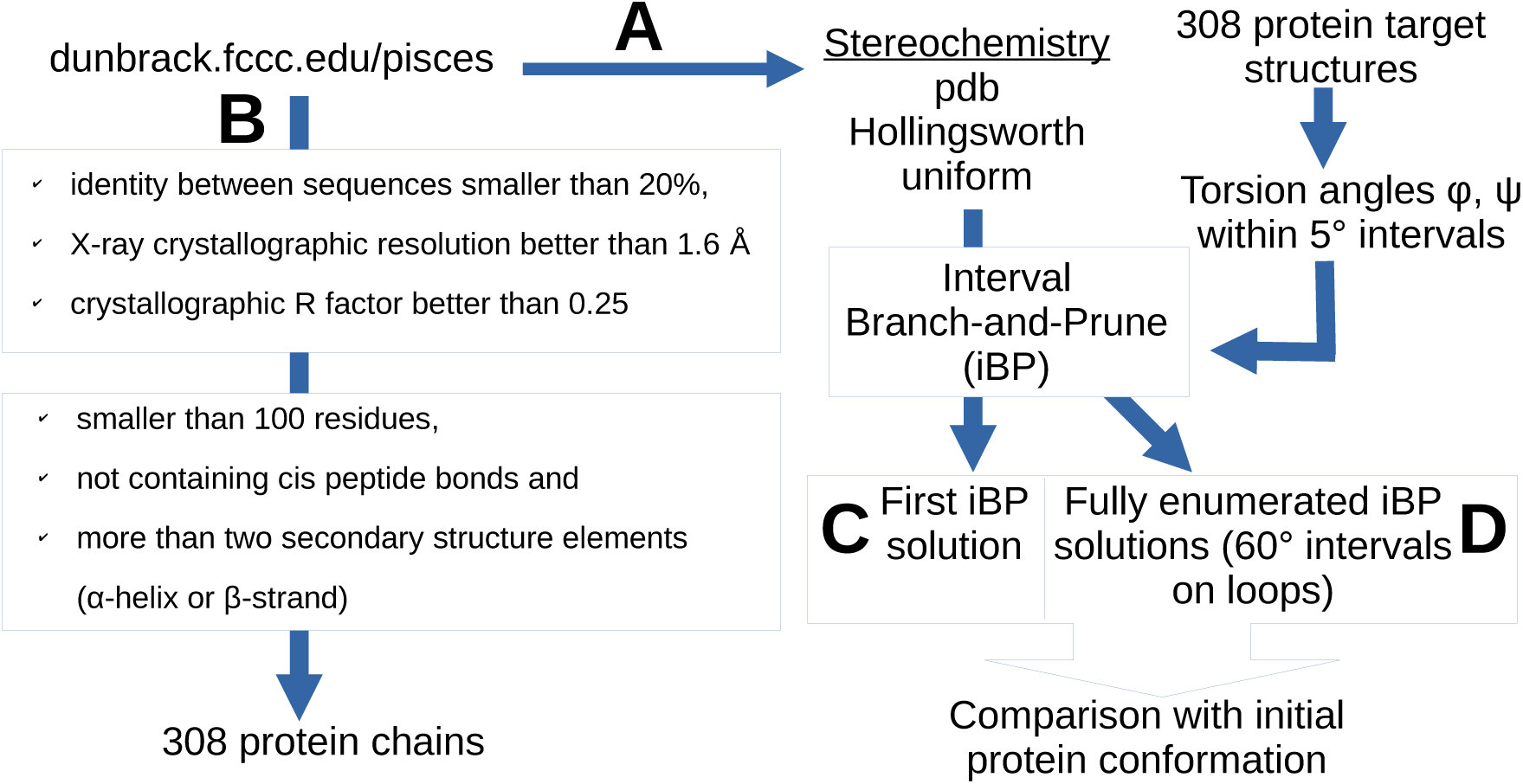
Flow-chart of the calculations. A. Obtaining the statistics of stereochemistry from a protein database. B. Generating a subset of 308 protein chains which will be the targets for iBP calculations. Using the torsion angle values measured on the protein target conformations along with different hypotheses on stereochemistry (Table 1), protein conformations were recalculated using iBP, selecting the first generated conformation (one-shot iBP run) (C) or enumerating all possible conformations (D).

**Table 1:** Definitions of stereochemistry inputs.

| name | origin of<br>stereochemistry |
| --- | --- |
| pdb | initial conformation from the Protein Data Bank |
| Hollingsworth | averaged angle values from the Ramachandran regions<sup>33</sup> |
| uniform | stereochemistry parameters from Engh and Huber<sup>34</sup> (Table S2) |

The present study shows that the efficiency of reconstructing the protein fold is very sensitive to the knowledge of stereochemistry variations, namely the variations of bond angle values between the heavy backbone atoms and of the torsion angles *ω*. We will show that these stereochemistry variations depend more on the position of the residues in the Ramachandran diagram than on the type of individual amino acid residues. From these statistics, different types of stereochemistry were investigated, in particular, the case where the stereochemistry parameters are averaged on the regions of the Ramachandran diagram defined by Hollingsworth et al^33^ and that we will denote in the following by *Hollingsworth stereochemistry*. Two other stereochemistry types we analyze are the uniform one, in which the parameters are taken from Engh and Huber,^34^ and the pdb one in which the stereochemistry parameters are extracted from each studied PDB entry. Using the Hollingsworth stereochemistry, the exact knowledge of the *ω* backbone angles allowed us to recover most of the protein folds. Even a discretized knowledge of *ω* allowed us to achieve decent reconstruction levels. The enumeration of conformations using the iBP approach improves the fold reconstruction whenever uniform stereochemistry is used. The calculations performed here have been summarized in the flow chart described in Figure 1. Looking at the origin of *ω* variability, some connection with the position of rare DNA codons was emphasized, in agreement with recent literature results.^35–37^

## Materials and Methods

### Data and Software Availability

The list of PDB entries of X-ray crystallographic structures with identity between sequences smaller than 20%, resolution better than 1.6 Å and R factor better than 0.25, has been downloaded from the server dunbrack.fccc.edu/pisces^38^ providing 3757 protein chains. From this list, 308 protein chains were selected, smaller than 100 residues, not containing *cis* peptide bonds, and for which more than two secondary structure elements (*α*-helix or *β*-strand) are present (Table S1 and Figure 1A).

The proteins forming the database display a size mostly in the range of 60-90 residues, with a smaller number of proteins containing 20 to 60 residues (Figure S1). The percentage of *α* helices is uniformly distributed among the proteins, whereas the *β*-strand and the loops display more concentrated distributions in the 0-20% for *β* strands and 20-40% for loops.

The version of iBP^28^ modified to handle variable stereochemistry is available at: github. com/tmalliavin/ibp-ng-fullchain. For the other software, not developed by the authors, the literature references are given.

### interval Branch-and-Prune approach

The interval Branch-and-Prune approach (iBP) algorithm was initially proposed by Mucherino and coworkers^25,26,28,39–42^ to enumerate the conformations of proteins verifying sets of distance constraints. The space of all possible protein structures is described as a tree and the available geometric information permits tree branching and pruning. Each time a branch of the tree is pruned, the iBP calculation is stopped and resumed at the previous positioned atom. This branch-and-prune description of the problem makes possible a discrete enumeration of solutions, and consequently strongly contrasts with most of the optimization approaches usually employed for the determination of biomolecular structure.

If not otherwise stated, the conformations of the proteins have been recalculated using one-shot iBP runs, in which the run was stopped after producing the first solution. The branching part was performed on *ϕ* and *ψ* torsion angles using intervals of 5*^◦^* centered around the true *ϕ* and *ψ* values. The torsion angles are converted into distance intervals, which are discretized with a maximum of four branches separated by at least 0.1 Å, which defines the discretization factor *ɛ*.

The *ω* values of the torsion angle of peptidic planes were used as pruning restraints as well as the *χ*_1_ torsion angle defining L amino acid residues. A last pruning restraint is related to all interatomic distances which should be larger than the sum of van der Waals radii of the atoms, using a scale factor of *ρ* = 0.8 on the radii if not otherwise stated. This approach is reminiscent of the reduction of the van der Waals interactions during the simulated annealing procedure in NMR structure calculation. ^43^

### Variations of stereochemistry during iBP calculations

Several definitions of protein stereochemistry focusing on the backbone bond angles and *ω* torsion angles were used as inputs for the calculations. Uniform stereochemistry was defined using the values from the force field PARALLHDG (version 5.3)^34^ (Table S2). Two variations of the stereochemistry are explored: (i) pdb stereochemistry in which the bond angles and *ω* torsion angle were extracted from the PDB conformation of the considered protein, (ii) Hollingsworth stereochemistry in which the bond angles and the *ω* torsion angle are taken as the average stereochemistry values calculated from the regions of the Ramachandran diagram defined by Hollingsworth et al from the analysis of high-resolution X-ray crystallographic structures.^33^ The correspondence between the regions displayed in Figure S1 and the definition of Ref^44^ is given in Table S3.

For pdb stereochemistry, each protein residue is defined by a 3-letter name, the alphabetic order of the names coding for the positions of the residues in the primary sequence, the first residue being AAA, the second one AAB, and so on. The topology files in CNS format^45^ were modified by using this residue code to define the amino acid residues along the primary sequence as well as the different atom types for each residue. Using these atom types, and the stereochemistry values in the PDB structure, values of bond lengths and angles are then generated for each residue along the sequence and stored in the CNS parameter file. This allows us to take into account any possible variations of protein stereochemistry (pdb or Hollingsworth) along the protein sequence.

### Analysis of obtained conformations

The analysis of protein conformations obtained with iBP has been performed using the MDAnalysis package^46^ and STRIDE.^47^ Sidechains were added to the protein backbone using the Relax procedure^48^ of Rosetta^49^ for the refinement of a one-shot iBP run, in the case of uniform and Hollingsworth stereochemistry. During the Relax procedure, 10 conformations were generated and the procedure was repeated 5 times.

## Results

### Analysis of protein stereochemistry

The stereochemistry of the 3757 protein chains downloaded from the server dunbrack.fccc. edu/pisces^38^ has been analyzed (Figure 1) by calculating the average values of the backbone bond angles N*−*C*_α_−*C, C*_α_−*C*−*N, C*−*N*−*C*_α_*, C*_α_−*C*−*O, and O*−*C*−*N (Figure 2). The negative torsion angles *ω* were shifted by 360*^◦^* in order to obtain *ω* value variations around 180*^◦^*. The averaged and standard deviation values of these bond and torsion angles are plotted according to the type of amino acid (Figure 3, left column) and to the region of the Ramachandran diagram defined by the backbone torsion angles *ϕ* and *ψ* (Figure 3, right column). The Ramachandran regions were taken from the definition given in the work of Hollingsworth et al^33^ (Figure S2).

**Figure 2:**
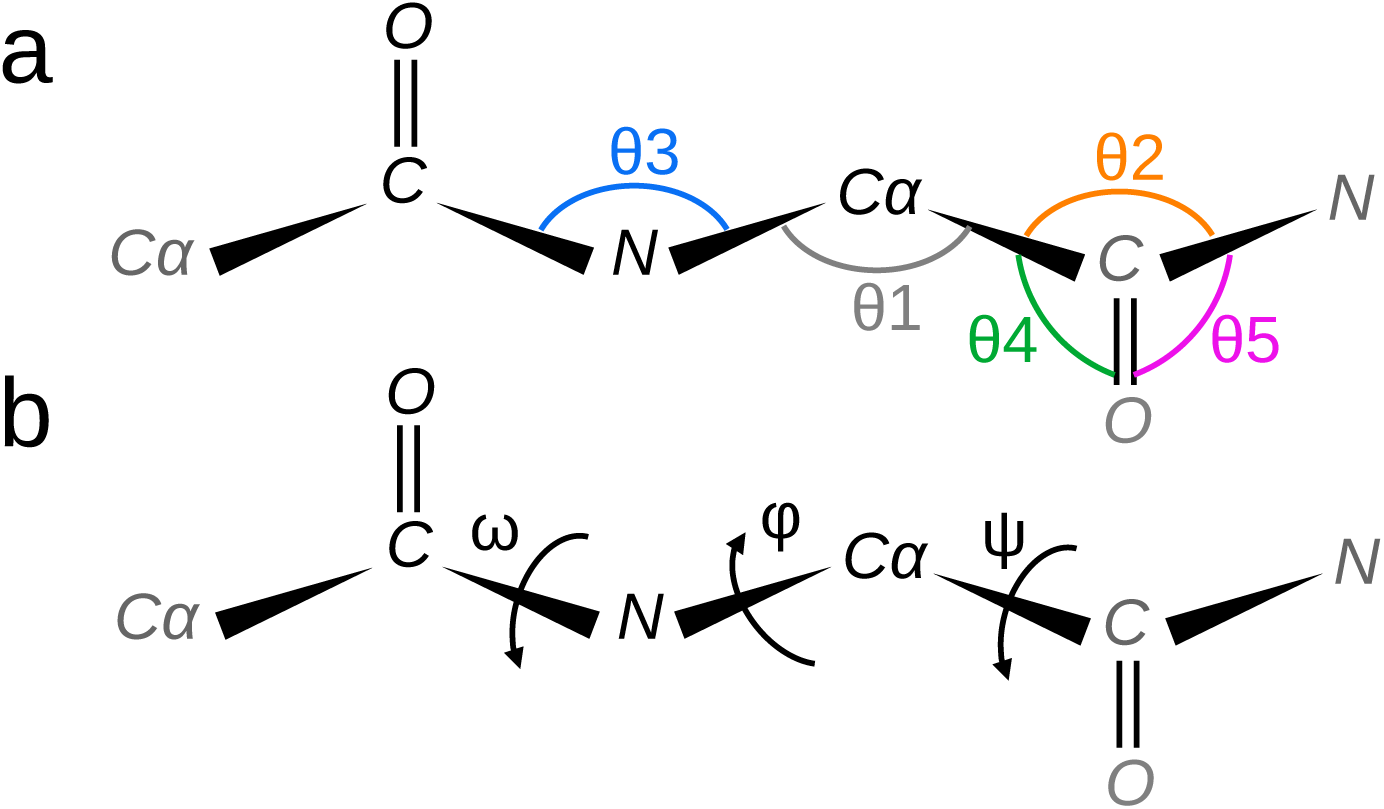
Scheme of the succession of protein backbone heavy atoms N, C*_α_*, C, and O along with definitions of angle parameters. a. Bond angles are: N*−*C*_α_−*C (*θ*_1_, grey), C*_α_−*C*−*N (*θ*_2_, orange) and C*−*N*−*C*_α_* (*θ*_3_, blue), C*_α_−*C*−*O (*θ*_4_, green) and O*−*C*−*N (*θ*_5_, magenta). b. The backbone torsion angles *ϕ*, *ψ*, and *ω* are indicated by circular arrows.

**Figure 3:**
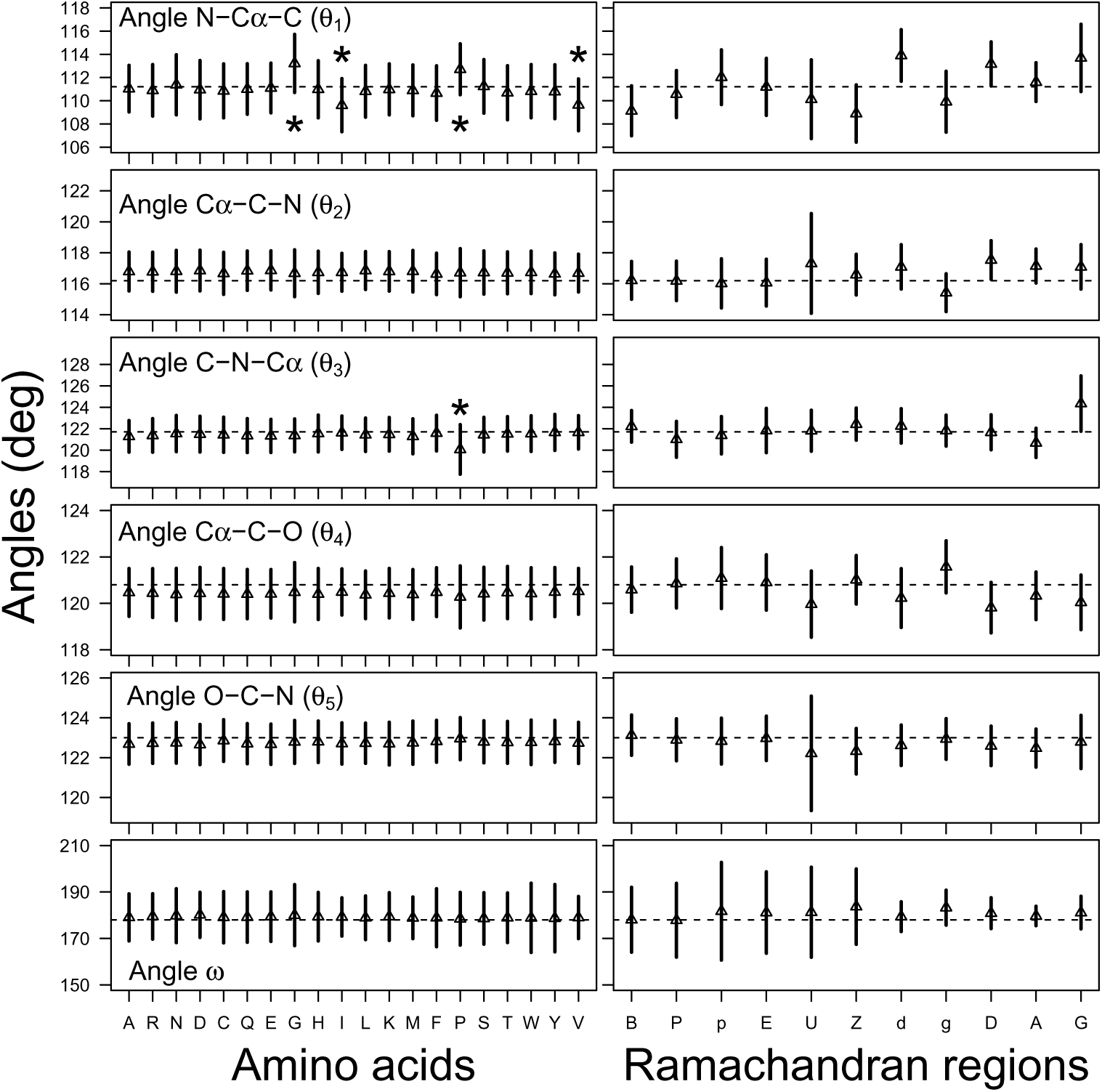
Average and standard deviation values calculated on the bond angles and *ω* dihedral angle, defining the stereochemistry of protein backbone. The bond angles labels are the same than those displayed on Figure 2. The regions of the Ramachandran diagram were taken from Ref^33^ and are displayed in Figure S2. The dashed lines correspond to the angle values in the parameter set of Engh and Huber^34^ (Table S2). Asterisk indicate the most variable bond angles along the amino acid type.

Almost all average angle and standard deviation values display flat profiles along the type of amino acid (Figure 3, left column). The standard deviations for *ω* angles display slight variations among the amino acids, especially for Glycine, Tryptophan, and Tyrosine (Table S4). Unsurprisingly, the dashed line indicating the Engh and Huber^34^ values is close to the average values of angles. The bond angles C*_α_−*C*−*N and C*−*N*−*C*_α_* display the smallest standard deviations, whereas the bond angles N*−*C*_α_−*C and C*_α_−*C*−*O display the largest ones. The averaged values of the angle C*−*N*−*C*_α_* display one outlier for Proline residues, with a shift of around 2*^◦^*. The averaged values of the angle N*−*C*_α_−*C display four outliers, all shifted by around 2*^◦^*: two are shifted towards larger values for amino acids Glycine and Proline, and two are shifted towards smaller values for amino acids Isoleucine and Valine. The outliers positions of Isoleucine and Valine have been recently observed^50^ for the propensity scales of the Ramachandran regions. In addition, Proline and Glycines have been known for decades to influence local geometry. ^51,52^

Interestingly, the profiles along the Hollingsworth regions (Figure 3, right column) are much more variable for the average as well as for the standard deviation values (Table S4). Among the bond angles, the angle N*−*C*_α_−*C and C*_α_−*C*−*O display the most variable profiles. The large variability of these angles may arise from the involvement of the atoms N and O into hydrogen bond network stabilizing the protein secondary structures.

The angle variability depends on the Ramachandran regions. The region A (Figure S2, red), corresponding to the regular *α* helix, produces angles close to the Engh and Huber values, with the smallest standard deviations. The region D (Figure S2, green), corresponding to the 3-10 helix, displays standard deviations similar to the region A, but average values shifted to upper values for bond angles C*_α_−*C*−*N and N*−*C*_α_−*C and to lower values for bond angle C*_α_−*C*−*O. The region B, corresponding to the regular *β* strand (Figure S2, blue), and P, corresponding to the polyproline region (Figure S2, brown), displays also average values mostly close to the Engh and Huber values except for the angle N*−*C*_α_−*C, but the standard deviations are larger, especially for the angle *ω*. The regions g, Z, located between the *α* and *β* regions of the Ramachandran diagram, the regions G, d, p, located in the loop region of positive *ϕ* value, and the region E, all display large standard deviations and shifted average values.

The right column of Figure 3 permits the definition of protein stereochemistry depending on the Ramachandran region by averaging over each Hollingsworth region (Figure S2), the values of bond angles and *ω* angles. Due to the profile variations of Figure 3, one may expect that this Hollingsworth stereochemistry will be more variable than a stereochemistry based on the amino-acid type. This is not surprising as the amino acid type is defined by the sidechains which are more far apart from the backbone than the *ϕ* and *ψ* torsion angles. In the following, the protein stereochemistry will be modeled as uniform ie. uniquely defined from the atom type, following the measurements of Engh and Huber^34^ (Table S2), as Hollingsworth with averaged values determined from the *ϕ*, *ψ* torsion angles (Figure 3 and Table S4), and as a pdb, using angles measured on the PDB structure of the considered protein.

### Effect of stereochemistry on the protein conformations generated with iBP

Several experiments were performed on the database of proteins to reconstruct the conformations using iBP with the previously chosen types of stereochemistry. First, one-shot runs were realized with calculations stopping after obtaining the first conformation (Figure 1C) and intervals of 5*^◦^* for *ϕ* and *ψ* angles. Then, the protein targets were submitted to a full exploration of the tree, using narrow intervals (5*^◦^*) of *ϕ* and *ψ* in the secondary structure elements, and larger intervals (40*^◦^*) in the connecting loops (Figure 1D).

Figure 4 displays the distributions of the root-mean-square deviation (RMSD, Å) between the atomic coordinates of the iBP solution and of the initial PDB conformation for the oneshot runs using the definitions of stereochemistry described in Table 1. In the present work, the RMSD values were calculated on the heavy atoms of the protein backbone. In the case where the stereochemistry is defined from the initial PDB conformation (pdb stereochemistry in Table 1), coordinate RMSD around 0.5 Å are observed (Figure 4a). This provides a floor value for the maximum possible precision which can be obtained using the discretization of the *ϕ* and *ψ* intervals in iBP. It is interesting to note that if the bond lengths are also taken variable from the PDB conformation, the same distribution of RMSD values is obtained (data not shown). The bond length variations have thus much less influence on the variations of conformations obtained by iBP than the bond angle variations.

**Figure 4:**
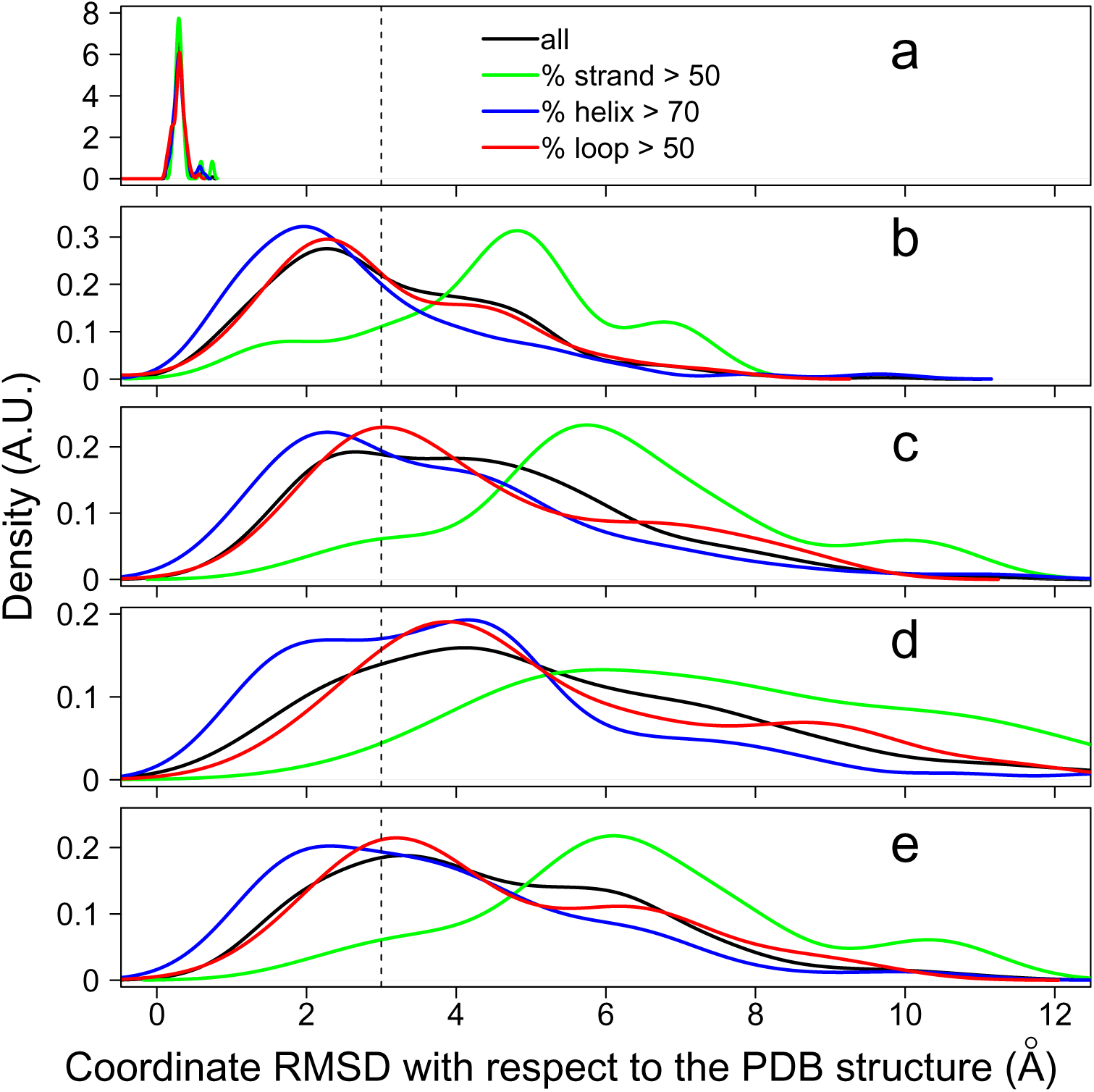
Distribution of the root-mean-square deviation (RMSD, Å) of atomic coordinates between the initial PDB conformation and the conformation reconstructed using iBP. The following stereochemistry inputs (Table 1) were used: (a) pdb stereochemistry taken from the PDB input, (b) pdb stereochemistry with *ω* values of 178 deg, (c) Hollingsworth stereochemistry with bond and *ω* angles averaged along the Hollingsworth regions (Figure S2), (d) Hollingsworth stereochemistry with *ω* values of 178 deg, (e) uniform stereochemistry^34^ (Table S2). The vertical dashed line indicated the RMSD value of 3 Å. The curves are colored depending on the percentage of residues belonging to *α*-helices, to *β*-strands, or to loops as described in the legend. The secondary structures were determined using STRIDE.^47^

As soon as the *ω* angle is set to 178*^◦^* while getting the other parameter values from pdb stereochemistry (Figure 4b), the RMSD distributions are switched towards much larger values, up to 10-12 Å. Such behavior is also observed for Hollingsworth (Figure 4c,d) or for uniform (Figure 4e) stereochemistry. Interestingly, quite different RMSD distributions are observed according to the type of secondary structures. The shift is smaller for proteins folded mostly as *α* helices (blue curves) or mostly as loops (red curves), producing an RMSD value smaller than 3 Å for at least half of the structures. By contrast, the structures containing mostly *β* strands (green curves) display distributions centered at RMSD values between 5 and 6 Å.

The coordinate RMSD values are known to display some limitations for precisely measuring the accuracy of a protein structure prediction.^53^ Thus, the TM score distribution^54^ have been calculated (Figure S3) using the software code downloaded from zhanggroup. org/TM-score. For pdb stereochemistry (Figure S3a), the TM scores are close to the optimal value of 1. Similarly to Figure 4, the TM-score values are getting worse if *ω* values of 178*^◦^* are used (Figure S3b,d) or in the case of uniform stereochemistry (Figure S3e). Most of the calculated conformations display TM-scores larger than 0.5, if the bond angles are taken from the PDB entry and the *ω* values are set equal to 178*^◦^* (Figure S3b).

An interesting difference between the TM score and RMSD is observed for Hollingsworth stereochemistry. Indeed, the TM scores are worse for Hollingsworth stereochemistry (Figure S3c,d) than for any other calculation, whereas the RMSD values are similar between Hollingsworth (Figure 4c,d) and uniform (Figure 4e) stereochemistry. This is probably due to a distortion in interatomic distance distribution, produced by the use of bond and *ω* angles averaged on Hollingsworth regions. Indeed, this distortion deteriorates the TM score value as the distance distribution is the main ingredient of the score. Additional distance distortions may arise from the use of the van der Waals scaling of *ρ* = 0.8 used during the iBP calculation to avoid pruning of conformations.

To investigate more precisely the relationship between stereochemistry variations and the efficiency in conformer generation, the global variation of bond angles along a structure has been calculated as:

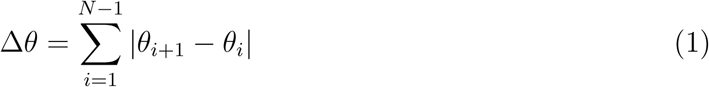

where *| · |* stands for the absolute value, and *N* is the number of residues with residue number indexed from 1 to *N*. A similar global variation for the torsion angle *ω* was defined as:

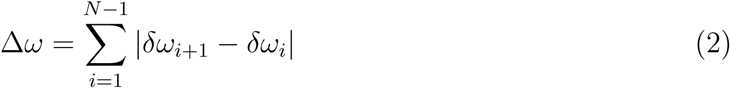

where:

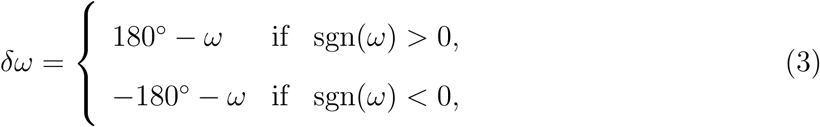

with sgn(*·*) being the sign function, i.e., sgn(*x*) = 1 if *x >* 0 or sgn(*x*) = *−*1 if *x <* 0.

The Δ*θ* values calculated on bond angles C*−*N*−*C*_α_*, N*−*C*_α_−*C and C*_α_−*C*−*N and the Δ*ω* values calculated on torsion angle *ω* were compared to the coordinate RMSD values between the iBP and initial conformations (Figure S4). The global variations and the coordinate RMSD display an obvious correlation which is also driven by the length of the protein chains. In agreement with Figure 3, the largest global variations are obtained for Δ*ω* (blue points) and Δ*θ* of the N*−*C*_α_−*C bond angle (green points).

For the calculations performed using: (i) Hollingsworth stereochemistry (Figure 4c) and (ii) uniform stereochemistry (Figure 4e), the protocol Relax^48^ of Rosetta^49^ was applied on the iBP outputs, to add the residue sidechains. The minimal RMSD value with respect to the initial PDB conformations (Figure S5a,b) shifts towards smaller values which is the sign of a conformation drift towards the correct solution. Indeed, the comparison of RMSD distribution with Hollingsworth (Figure S5a versus Figure 4c) and uniform (Figure S5b versus Figure 4e) stereochemistry reveals a shift of 1-2 Å and even of 4 Å for the mostly *β* folded proteins (green curve). The Rosetta scores have been also plotted along the coordinate RMSD and display a similar variation towards more negative values for smaller RMSD values (Figure S5c,d).

The iBP procedure presented here for reconstructing a protein complete fold could also have an application for the reconstruction of missing parts of a given protein structure. To evaluate this approach, the sub-chains for which coordinate RMSD to initial protein structure was smaller than 2.5 Å were extracted and their lengths are plotted as the percentage of the length of the full chain (Figure S6a,b,c) as well as numbers of residues (Figure S6d,e,f). The distribution of the percentages (Figure S6a,b,c) agrees with the distribution of RMSD values (Figure 4), with percentages close to 100% when RMSD values close to 0.5 Å are observed. As soon as the stereochemistry becomes less variable, the distributions of percentages become wider, but display very similar shapes in all runs, with two maxima located around 50% and 90% (Figure S6b,c). The distribution of the numbers of residues are all larger than 30 residues and are mostly distributed in the range of 20-60 amino acids. These values compare well with the results of the literature.^55^ In addition, similar distributions are observed for the different types of secondary structures in protein folds. These results are quite encouraging in the perspective of reconstructing non visible regions in protein structures.

In the presence of Hollingsworth stereochemistry, the effect of different input *ω* values on the reconstruction of protein folds was analyzed (Figure 5). If exact values are known for the *ω* torsion angles (Figure 5a), the majority of structures containing mostly *α* helices or loops display RMSD values smaller than 3 Å, corresponding to a good reconstruction of the protein fold. On the other hand, the structures containing mostly *β* strands display a shift in RMSD values, but their RMSD is still mostly smaller than 3 Å. Thus, knowing the exact values for torsion angles *ω* is essential for building the protein structure from local information.

**Figure 5:**
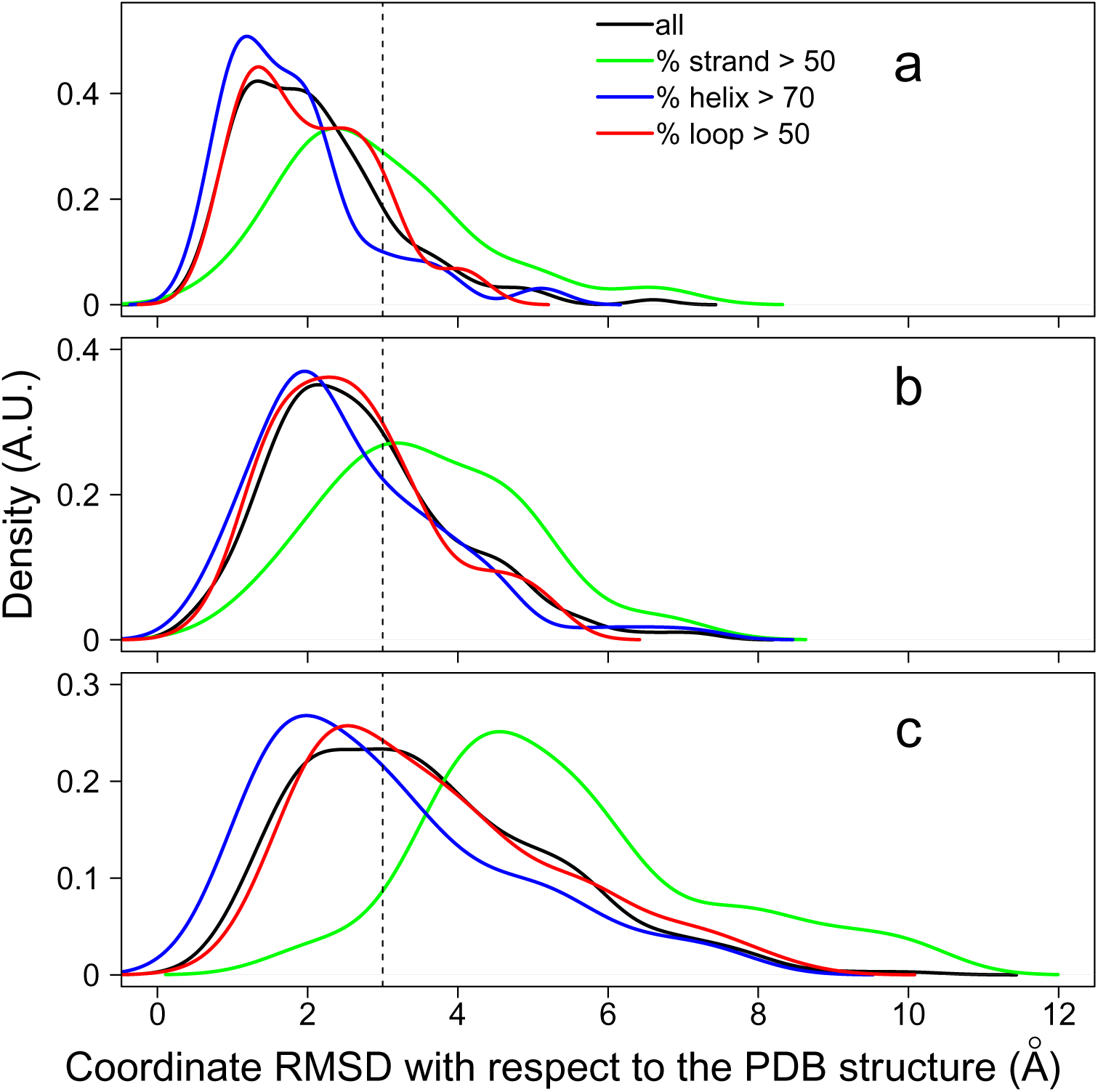
Distribution of the root-mean-square deviation (RMSD, Å) of atomic coordinates between the initial PDB conformation and the conformation reconstructed using iBP with the Hollingsworth stereochemistry for bond angles along with various definitions of the *ω* angles: (a) *ω* values taken from the PDB initial conformation, (b) discretization of *ω* values among four classes (Eq 4), (c) discretization of *ω* to 178*^◦^*sgn(*ω*), where sgn(*ω*) is the sign of *ω* in the initial PDB structure.

Then, the effect of several discretizations of *ω* was tested on the reconstruction of protein structures. In that case, the *ω* continuous values are replaced by *ω_k_* values corresponding to different discretization classes *k*. In the first discretization, the absolute value of the parameter *δω* previously introduced in Eq 3 was used to define four classes of *ω* values:

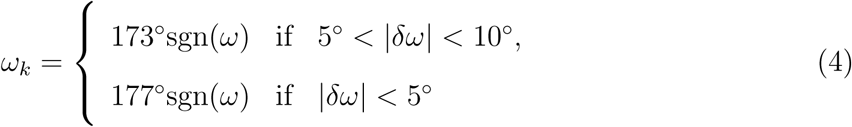

This discretization induces a shift in RMSD values (Figure 5b). The mostly *α* and loop structures are still correctly reconstructed, and half of the mostly *β*structures display RMSD values larger than 3 Å.

A more crude discretization is used where *ω_k_* is set equal to 178*^◦^*sgn(*ω*). This two-class discretization (Figure 5c) shifts the RMSD distribution to values larger than 3 Å for *β* structures, but about the two third of *α* and one-half of loop structures display RMSD values smaller than 3 Å. But, even this crude discretization allows us to obtain better RMSD values than those observed for uniform stereochemistry (Figure 4e). The effect of *ω* discretization on the fold reconstruction proves that classification approaches^56^ could be interesting for predicting protein conformations.

### Effect of the enumeration by iBP to the reconstruction of protein fold

The iBP approach has the advantage of allowing a systematic enumeration of all possible solutions. This enumerating scheme was thus used here to improve the coordinate RMSD of solutions with respect to the initial PDB structure. The inputs of the iBP runs were intervals around the *ϕ* and *ψ* angles with intervals widths of 5*^◦^* in *α* helices and *β* strands, and of 40*^◦^* in other protein regions. The number of branches is 4.

A disadvantage of the iBP approach is that execution can take a very long time and ultimately prune all solutions. In order to quickly determine input values avoiding the full pruning of solutions, short iBP runs were launched with an upper limit of 2 minutes, varying systematically the values of the discretization factors *ɛ* and of the van der Waals scaling *ρ*. Two stereochemistry inputs were used: uniform and Hollingsworth stereochemistry. A conformation was stored only if the coordinate RMSD between the newly generated and the previous solution was smaller than 3.5 Å.

The number of accepted solutions is mostly around 10^4^ and increases up to 10^5^ (Figure S7a). Around 20% of the calculations display no solutions. The number of solutions rejected because of the RMSD criterion (Figure S7b) is in the range of 10^6^-10^7^, much larger than the range of accepted solutions. An RMSD of 3.5 Å is thus quite discriminating for selecting solutions. The tree size is mostly in the range of 10^5^ to 10^10^ (Figure S7c). Interestingly, previous experiments realized with TAiBP showed^29^ that a tree size of about 10^9^ permits systematic enumeration of the tree solutions for protein fragments. The size of the trees, as well as the numbers of accepted and rejected solutions, display the same distribution for the Hollingsworth or the uniform stereochemistry. Similarly, the discretization factors *ɛ* vary uniformly in the range 0.15-0.17 (Figure S7d) for Hollingsworth (black curve) as well as for uniform (red curve) stereochemistry. By contrast, the van der Waals scaling *ρ* varies in the 0.2-0.6 range for Hollingsworth stereochemistry and in the 0.3-0.6 range for the uniform stereochemistry (Figure S7e). This shows that smaller *ρ* values were sometimes used to avoid pruning in the case of Hollingsworth stereochemistry and agrees with the worse TM score observed for the one-shot run with Hollingsworth stereochemistry (Figure S2).

Based on the fast exploration described above, input values for enumerating runs were selected using the following rules: (i) the largest possible van der Waals scaling *ρ* for maximizing the pruning by steric hindrance, (ii) the largest possible discretization factor *ɛ* for obtaining the smallest possible tree size to facilitate its full exploration. The corresponding trees were then completely parsed using iBP. During the enumeration, the number of asked conformations was set to 10^9^. All calculations produced a smaller number of conformations, which proves that the corresponding trees were fully explored. Tree sizes centered around 10^4^, discretization factors *ɛ* around 0.17, and van der Waals scaling factors *ρ* around 0.5 were used for these full runs (Figure S8). The discretization factor displays similar distributions for Hollingsworth and uniform stereochemistry. In contrast, the tree size and the van der Waals scaling factor *ρ* are slightly shifted towards higher values for Hollingsworth stereochemistry. Indeed, the larger tree observed for this stereochemistry requires greater van der Waals scaling to reduce the number of solutions by pruning.

The effect of the enumerating scheme for calculating structures was evaluated using the distribution of coordinate RMSD between iBP and PDB target conformations (Figure 6). For each processed protein, the smallest RMSD value between the iBP solution and the initial structure was selected and the corresponding RMSD distribution was compared to the corresponding RMSD distributions for the one-shot runs (Figure 4c,e). For both uniform (Figure 6a) and Hollingsworth (Figure 6b) stereochemistry, the use of enumeration induces a shift of the RMSD values towards smaller values. Interestingly, this shift is more pronounced in the case of uniform stereochemistry, as shown by the comparison of Figures 6b and 4e. Thus, using the enumeration of conformations potentially improves the efficiency of the fold reconstruction.

**Figure 6:**
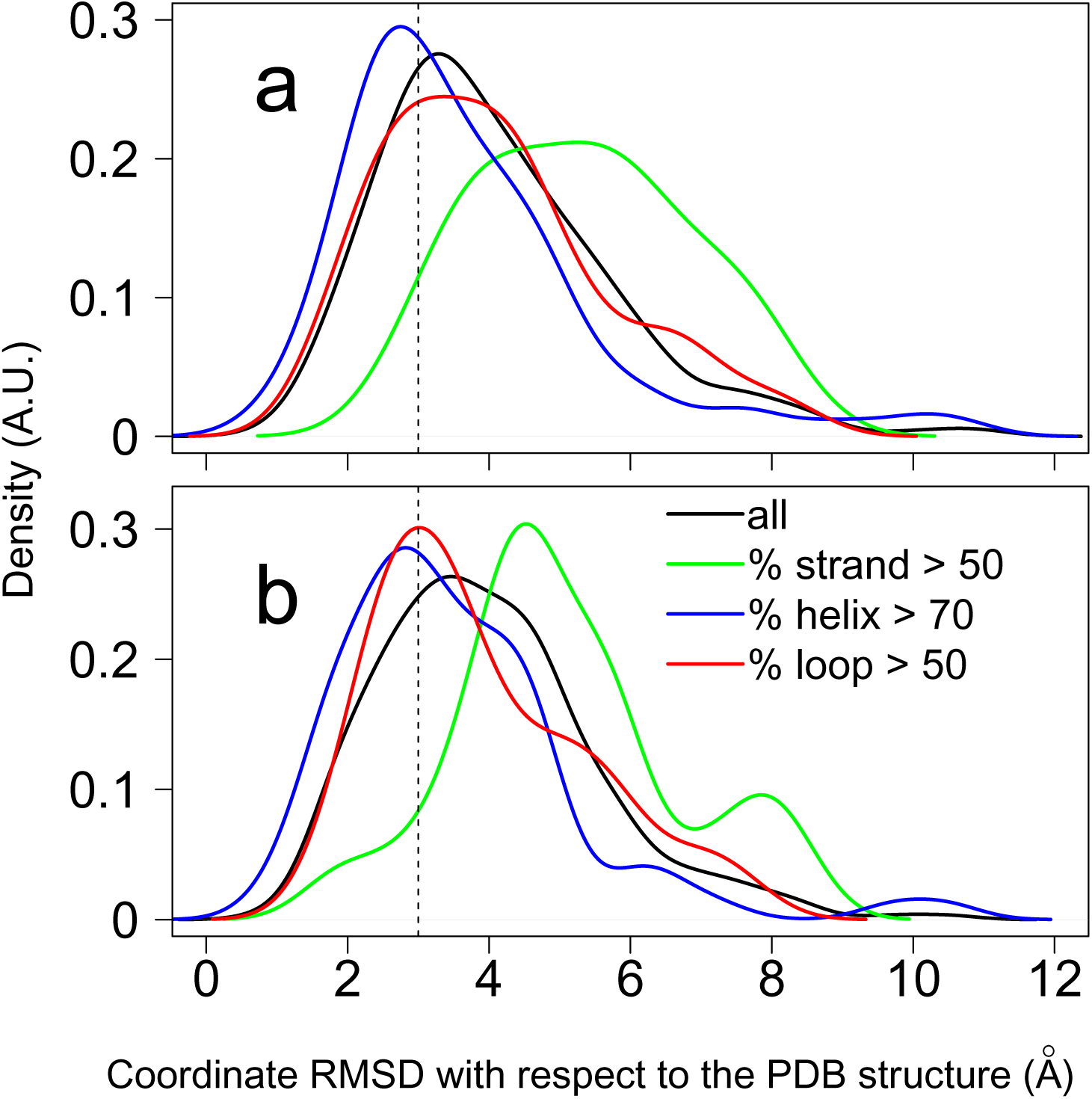
Distribution of the root-mean-square deviation (RMSD, Å) of atomic coordinates between the initial PDB conformation and the conformation reconstructed using enumerating iBP runs with Hollingsworth (a) or uniform (b) stereochemistry. The coordinate RMSD was taken as the smallest RMSD value obtained among all iBP solutions.

### A possible origin of the variability of stereochemistry

During the previous sections, the effect of variability of stereochemistry on the calculation of protein conformation based on local conformational restraints has been examined in various situations. In this section, we intend to investigate the relationship between the distribution of synonymous DNA codons and the variability of stereochemistry.

We first focused on the variability of bond angle values. The standard genetic code^57^ was used to determine the number of synonymous codons for each amino acid residue. The number of possible synonymous codons for each residue was summed along each of the 308 protein primary sequences to produce the cumulative number of synonym codons. Plotting the global variations Δ*θ* of the bond angles C*−*N*−*C*_α_*, N*−*C*_α_−*C and C*_α_−*C*−*N compared to this cumulative number (Figure S9) reveals a correlation between the stereochemistry variation and the number of synonymous codons similar to those previously observed in Figure S4. As in Figure S4, the correlation is driven by the protein size. The 13 proteins from *E coli* and expressed in *E coli* for structure determination are marked with green crosses and display the same tendency as the whole set of proteins.

These 13 *E coli* proteins are drawn in cartoon and the residues displaying global variations Δ*θ* of bond angles larger than 6*^◦^* are drawn in licorice and colored in green (Figure S10). Most of these protein structures display a topology inducing interactions between secondary structure elements located apart in the protein sequence. Also, the residues with the largest local variation of bond angles are mostly located in loops or at the extremity of secondary structure elements. In several structures (1C4Q, 1GYX, 1Q5Y, 3CCD, 4MAK, 4Q2L), most variable residues are close to each other in the 3D structure, displaying even long-range physico-chemical interactions. The variations of bond angle stereochemistry can be thus related to the long-range interactions participating to the fold definition. The positions of variable residues in the loops might be related to the importance of loop conformations for orienting the protein backbone with the folded topology. In addition, the long-range interactions of some variable residues suggest a cooperative effect between bond angle variations arising during the protein folding.

In the second step, we focused on the relationship between the *ω* torsion angle variability and the individual corresponding DNA sequences. Among the 308 protein structures, the proteins issued from the organisms *Homo sapiens*, *Escherichia coli* and *Saccharomyces cerevisiae* were selected using the descriptor SOURCE: ORGANISM SCIENTIFIC. The relative codon usage observed in three organisms: *Escherichia coli*, *Saccharomyces cerevisiae* and *Homo sapiens* (Table 1 of Ref^58^) was used to extract the different numbers of synonymous codons for each amino-acid residue of these proteins. The PDB entries were then entered as a query to the European Nucleotide Archive (ENA) www.ebi.ac.uk/ena. The corresponding DNA sequences were programmatically downloaded and filtered to keep those corresponding to the considered protein chain in the PDB entry.

The DNA sequence codons were then analyzed using the statistics on codons from Ref ^58^ calculated on the organisms *Homo sapiens*, *Escherichia coli* and *Saccharomyces cerevisiae*. From each amino acid, the codons displaying statistics of presence smaller than the average presence of all codons coding for the amino acid were considered rare codons. Then, the *ω* angle values of all protein residues were analyzed (Figure 7) by calculating their average *µ* and standard deviation *σ*^2^ values on each considered protein sequence. The *ω* angle values were centered and normalized using *µ* and *σ*^2^, producing a global averaged *ω* value on each protein equal to zero. The *ω* values averaged on protein residues corresponding to rare codons, as well as to protein residues corresponding to neighbors or second-neighbors of rare codons were centered and normalized using the *µ* and *σ*^2^ values obtained for the full sequence of the corresponding PDB entry. The distributions of these centered and normalized *ω* values display slight shifts towards positive or negative values (Figure 7a). Looking at these distributions, the *ω*values are more apart from zero for residues corresponding to rare codons than for neighbor and second-neighbor residues (Figure 7a). All standard deviation values (Figure 7b) are centered around 1*^◦^*, similarly to the standard deviation values normalized on the whole primary sequence.

**Figure 7:**
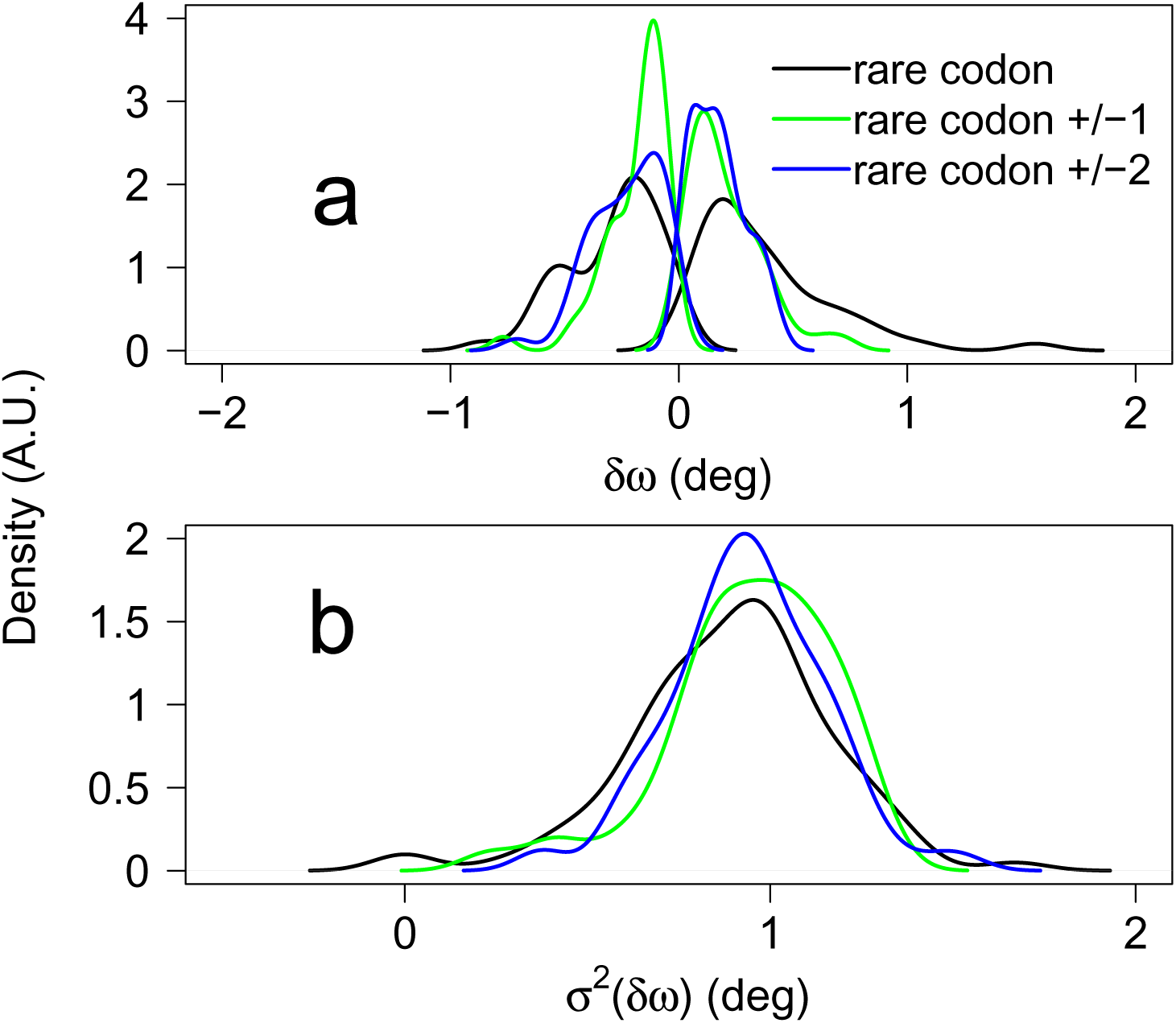
Distribution of the variations of *δω* (Eq 3) (a) and of the standard deviations *σ*^2^(*δω*) (b) for various positions in the protein sequences: at the residues for which the rare codons are observed (black curve), at the residues neighboring the rare codon (green curve) and at the residues second neighbor of the rare codon (blue codon).

The rare codons have been pointed out to be related to the kinetics of protein folding during the protein synthesis in the ribosome.^57,59,60^ In addition, recent bioinformatics analysis has established a relation between the genetic code and the protein structure.^35,36^ In that frame, the relationship put in evidence here between the variability of *ω* values, the rare codons, and the reconstruction of the protein structure connects the protein folding and the kinetics of protein synthesis in the ribosome.

In that respect, it is interesting to observe that mostly *β* folded proteins are specifically sensitive to the variability of *ω* values. Indeed, their folding requires intricate cooperation between the establishment of long-range interactions forming the *β* sheets. This may be related to the analysis of Figure S10 performed above.

The analyses performed here point out the importance of mRNA in the variability of stereochemistry in proteins. They complement the relationships put in evidence in the literature^35^ between the mRNA sequence and populations of *α* and *β* regions, as we have also shown here that the variations of stereochemistry are related to the Hollingsworth regions of the Ramachandran diagram (Figure 3).

## Discussion

The present work has been investigating the exclusive use of local conformational information, namely the values of the torsion angles *ϕ* and *ψ* for calculating protein conformations. The results obtained here were made possible in an essential way by the development of the interval Branch-and-Prune approach (iBP), ^40^ providing a framework for the systematic enumeration of conformations. The analyses performed here have put in evidence the essential impact of the variability in stereochemistry and represent, to the best of our knowledge, the first attempt to relate these stereochemistry aspects to the calculation and prediction of protein conformations.

The variations of stereochemistry are certainly influenced by the refinement protocols used for determining X-ray crystallographic structures, in which the application of longrange restraints can produce effects in variations of local stereochemistry, in a way that is not mastered in the details. During the last decades, the stereochemistry aspects have not been taken into account during the protein structure prediction thanks to the use of longrange distance/angle restraints. ^61^ On the other side, the use of long-range restraints might influence the appearance of stereochemistry outliers. The relative weights of the different types of information in the protein structure calculation should be further investigated for example using a Bayesian approach. ^62,63^

To alleviate the impact of variability, a Ramachandran-based definition of the bond angle stereochemistry, the Hollingsworth definition, has been proposed. The efficiency of this definition is improved with the use of enumeration during the iBP approach or by the knowledge of *ω* values. The combination of these aspects provides thus a way to overcome the variability problem for most of the protein structures examined here, especially in the case of *α* proteins.

The calculations performed here have been scored with respect to reference protein conformation, using coordinate RMSD and TM-score. In most of the calculations, TM-scores display better values than RMSD, in agreement with the general knowledge on this score.^54^ But, if Hollingsworth stereochemistry is used, better RMSD values are obtained than the TM-score values, probably because the deformation of local stereochemistry impacts the distribution of inter-atomic distances used in the TM-score calculations. Indeed, the TMscore was derived to correct the bias of coordinate RMSD on structures determined in the framework of uniform stereochemistry and should be adapted to the case of Hollingsworth stereochemistry.

Two approaches have been used to reduce the conformational drift produced by the lack of precision in the modeling of stereochemistry: the enumeration of conformation in the framework of iBP, and the Relax procedure^48^ of Rosetta.^49^ Both approaches permit to improve the results.

The analyses carried out here make it possible to propose that the origin of stereochemical variations could be linked to the information contained in the mRNA sequence. The finer investigation of this aspect is out of the scope of the present work but could provide a more integrated modeling of protein structure and folding.

## Acknowledgements

CNRS, Lorraine University, IRISA, and ANR PRCI multiBioStruct (ANR-19-CE45-0019) are acknowledged for funding. High Performance Computing resources were provided by the EXPLOR Centre at Lorraine University (2022CPMXX2687).

