## Supplementary Material for "Influence of stereochemistry in a local approach for calculating protein conformations"

September 16, 2024

[1] Laboratoire d'informatique de l'École Polytechnique, CNRS UMR 7161
[2] Institut de Recherche en Informatique et Systèmes Aléatoires, CNRS UMR 6074, University of Rennes
[3] Laboratoire de Physique et Chimie Théoriques (LPCT), University of Lorraine, Vandoeuvre-lès-Nancy, France,
[*] Corresponding author

September 16, 2024

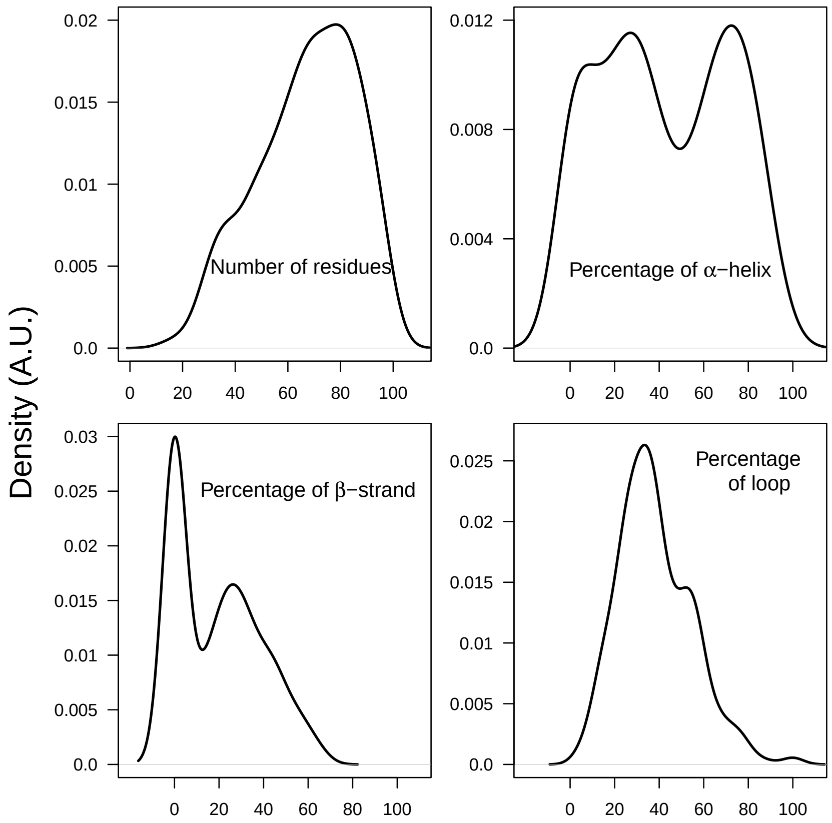


Figure S1: Distribution of number of residues and of percentages of secondary structures in the database of 308 proteins used for evaluating the iBP reconstruction.

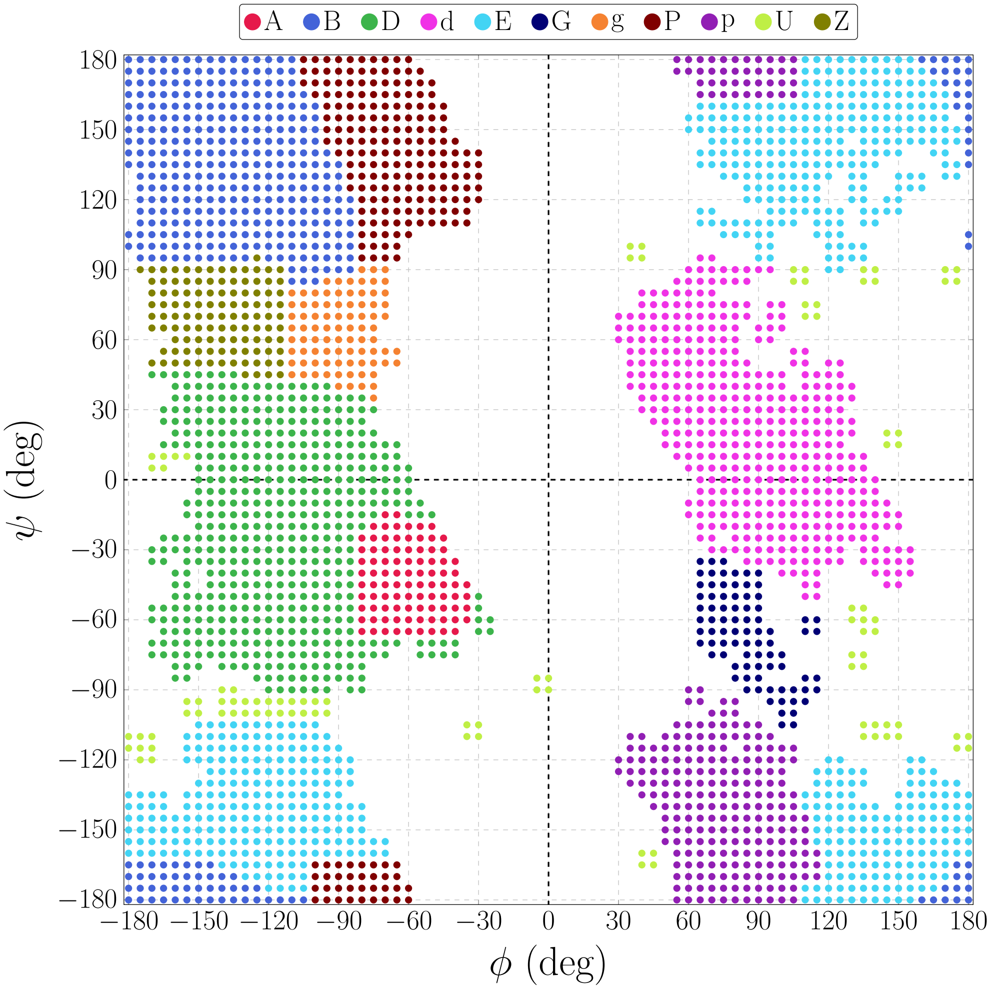


Figure S2: Ramachandran plot displaying the regions defined by Hollingsworth et al [1].

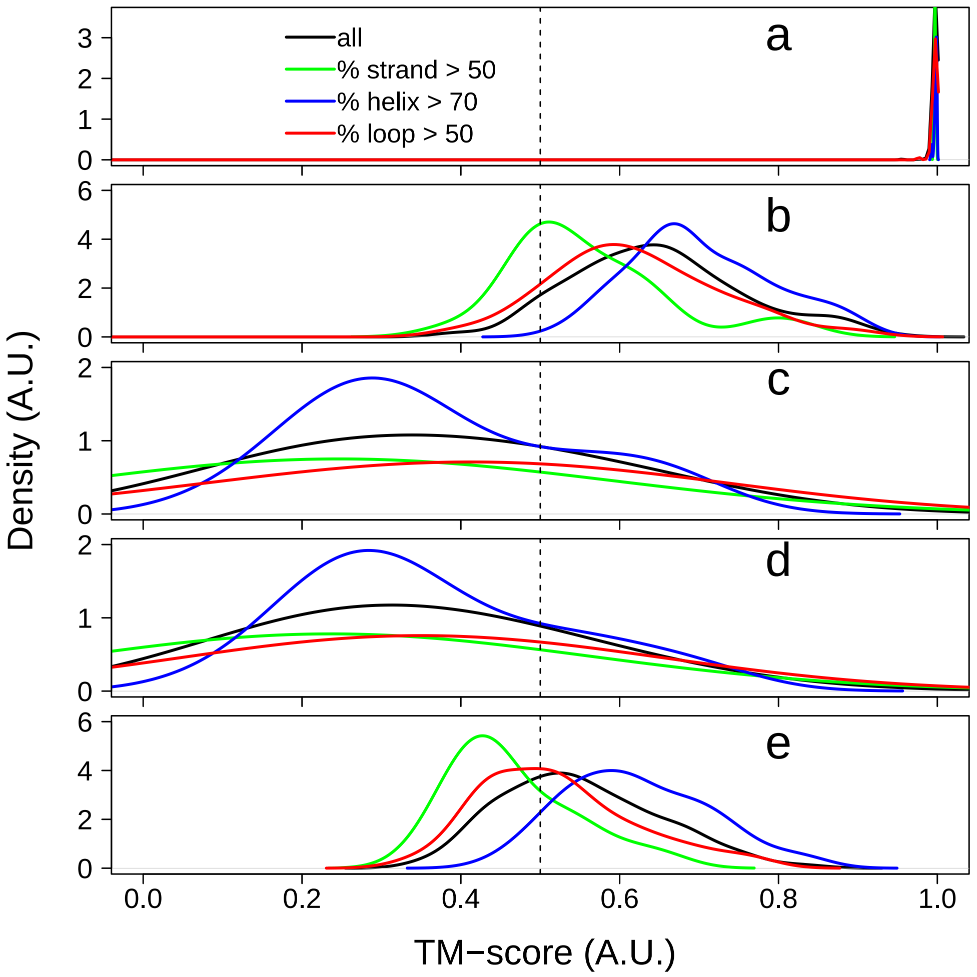


Figure S3: Distribution of the TM-score for the conformations reconstructed using the iBP approach corresponding to the runs shown in Figure 2 of the main text. The following stereochemistry inputs (Table 1 of the main text) were used: (a) pdb stereochemistry, (b) pdb stereochemistry with $\omega$ values of 178 deg, (c) Hollingsworth stereochemistry (Figure S3), (d) Hollingsworth stereochemistry with $\omega$ values of 178 deg, (e) uniform stereochemistry [2] (Table S2). The vertical dashed line indicates the TM-score value of 0.5. The curves are colored depending on the percentage of residues belonging to $\alpha$-helices, to $\beta$-strands, or to loops as described in the legend. The secondary structures were determined using STRIDE [3].

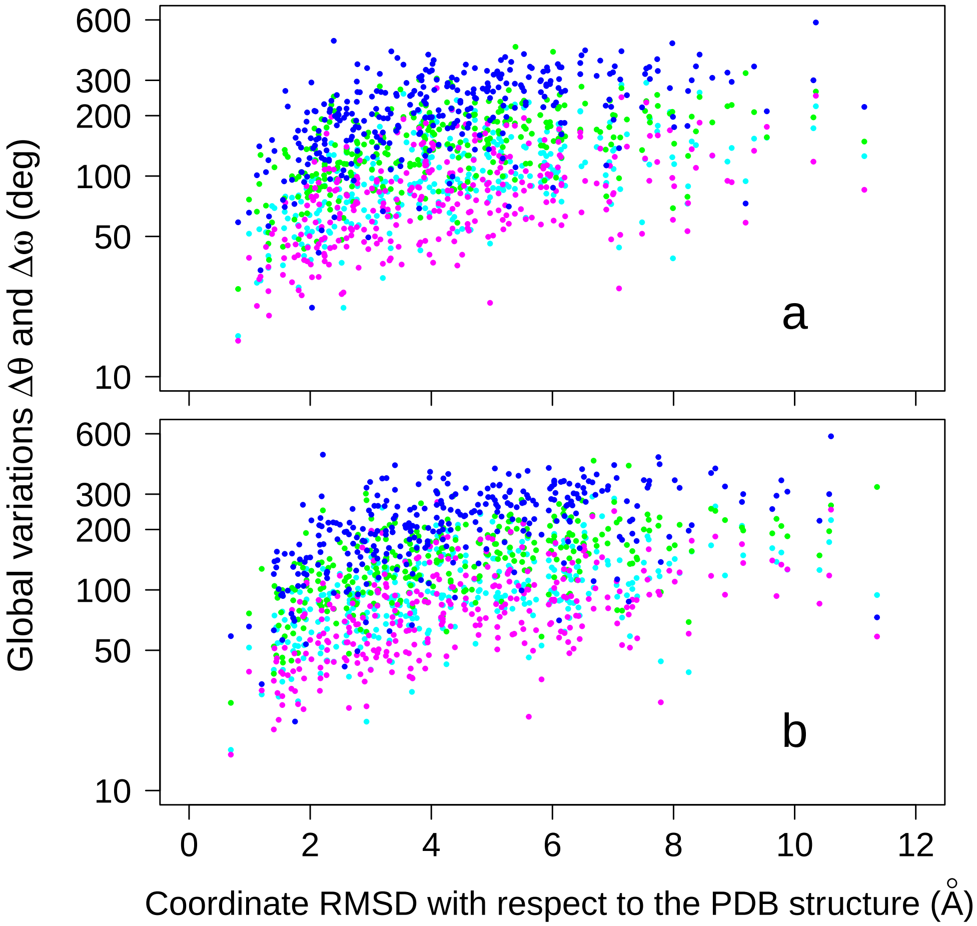


Figure S4: Global variations of bond angles ($\Delta\theta$, Eq. 1 of the main text) and of the torsion angle $\omega$ ($\Delta\omega$, Eq. 2 of the main text) plotted according to the coordinate RMSD with the protein target structure. The iBP calculations were performed using Hollingsworth (a) and uniform (b) stereochemistry. The analyzed stereochemistry parameters are the bond angles: $C{-}N{-}C_\alpha$ (cyan points), $N{-}C_\alpha{-}C$ (green points) $C_\alpha{-}C{-}N$ (magenta points) and the torsion angle $\omega$ (blue points).

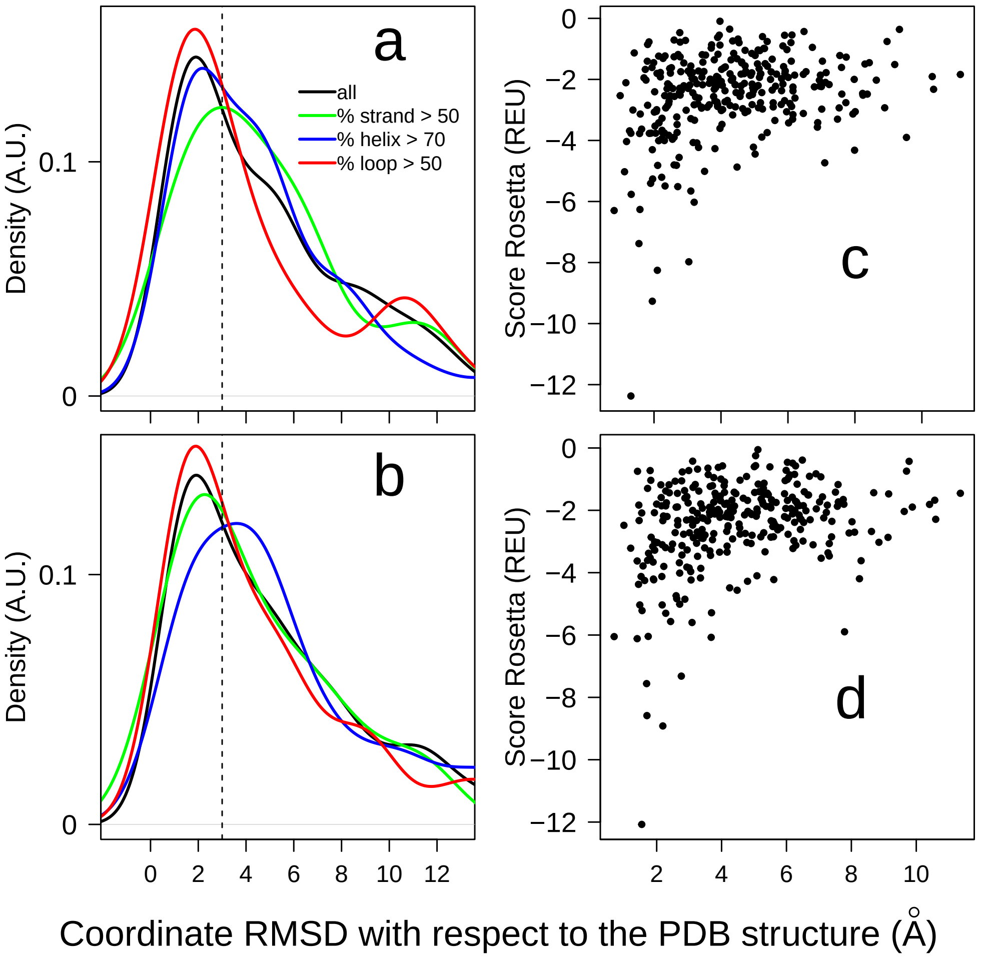


Figure S5: Effect of the sidechain addition to the results of iBP. The sidechains were added using the Relax procedure [4] of Rosetta [5]. Distribution of the coordinate RMSD to the target PDB conformation for conformations obtained with Hollingsworth (a) and uniform (b) stereochemistry. The conformations to which the sidechains were added are those for which the RMSD distributions are respectively plotted in Figures 4c and 4e of the main text. The vertical dashed line indicated the RMSD value of 3 Å. Rosetta scores plotted along the coordinate RMSD, obtained from Hollingsworth (c) and uniform (d) stereochemistry.

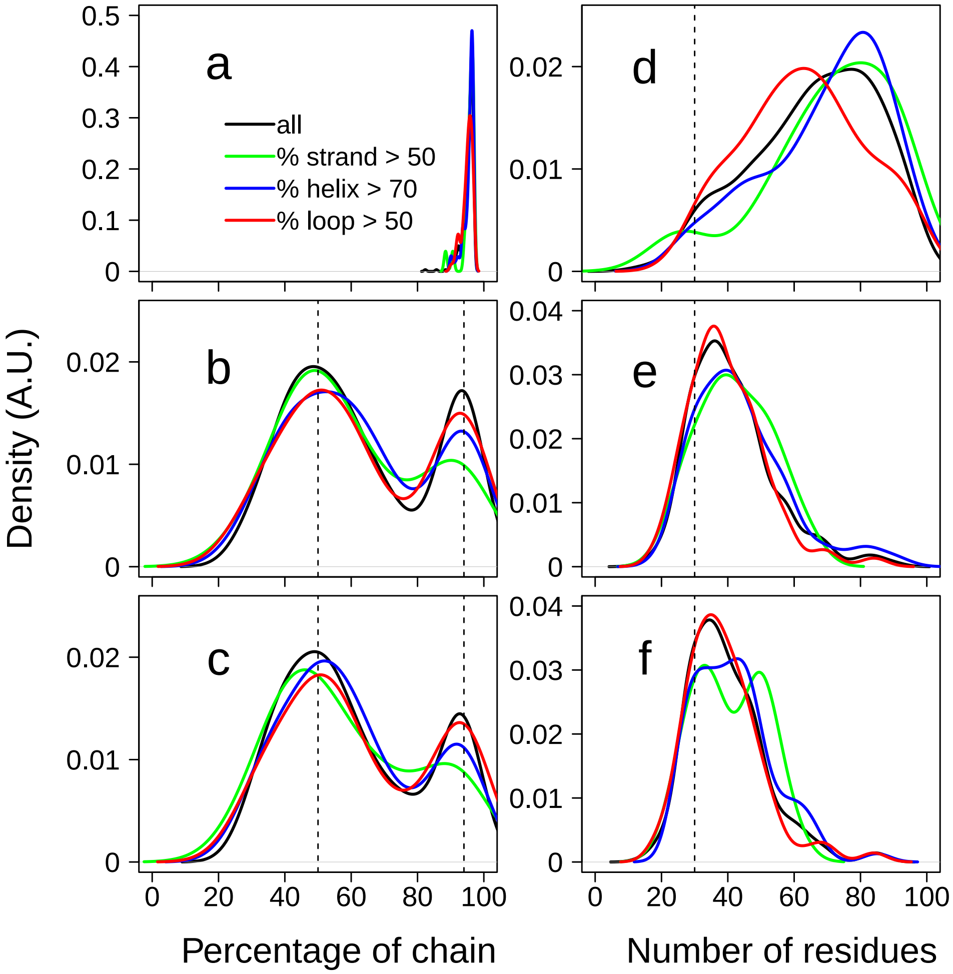


Figure S6: Distribution of the length of sub-chains reconstructed using iBP. The sub-chain lengths are expressed as percentages of the total length of the chain (a,b,c) or as the number of residues (d,e,f). The pdb (a,d), Hollingsworth (b,e), and uniform (c,f) stereochemistry inputs (Table 1 of the main text) were used. The curves are colored depending on the percentage of residues belonging to $\alpha$-helices, to $\beta$-strands, or to loops as described in the legend. The secondary structures were determined using STRIDE [3]. (b,c) The vertical dashed lines indicate the percentages of 50% and 94%. (d,e,f) The vertical dashed lines indicate the length of 30 residues.

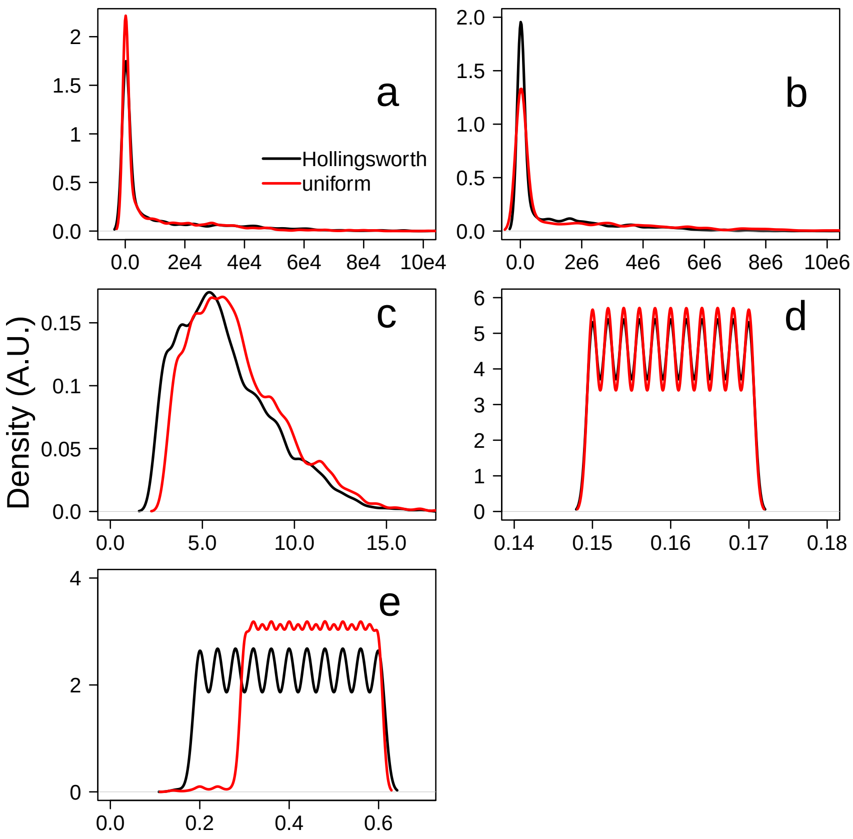


Figure S7: Distribution of the inputs values during the 2 minutes exploratory runs: (a) the number of accepted solutions, (b) the number of rejected solutions, (c) the decimal logarithm of the tree size, (d) the discretization factor $\epsilon$ and (e) the van der Waals scaling. For each parameter, the distribution curve is plotted for the Ramachandran (black curve) and uniform stereochemistry (red curve).

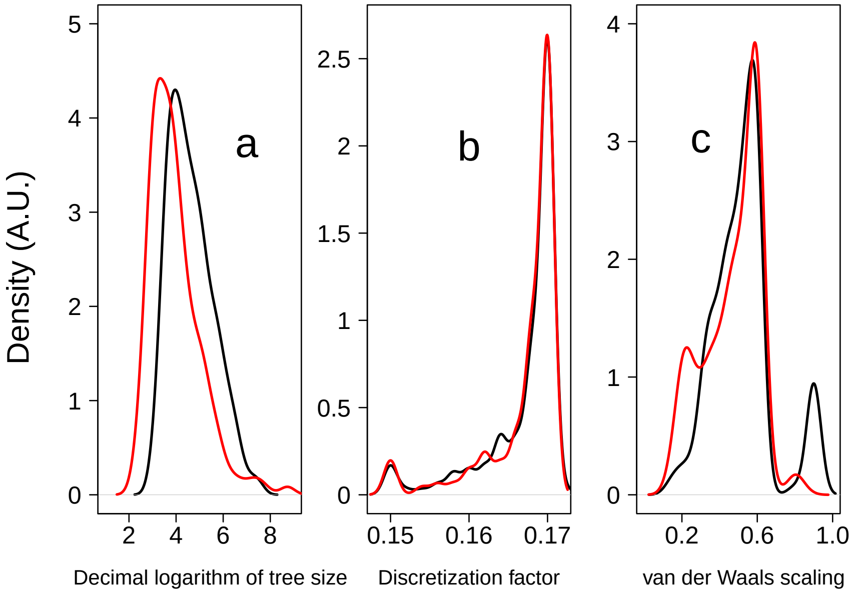


Figure S8: Distribution of the decimal logarithm of the tree size (a), of the discretization factor (b), and of the van der Waals scaling (c) during the enumerating iBP runs performed with Hollingsworth (black) and uniform (red) stereochemistry.

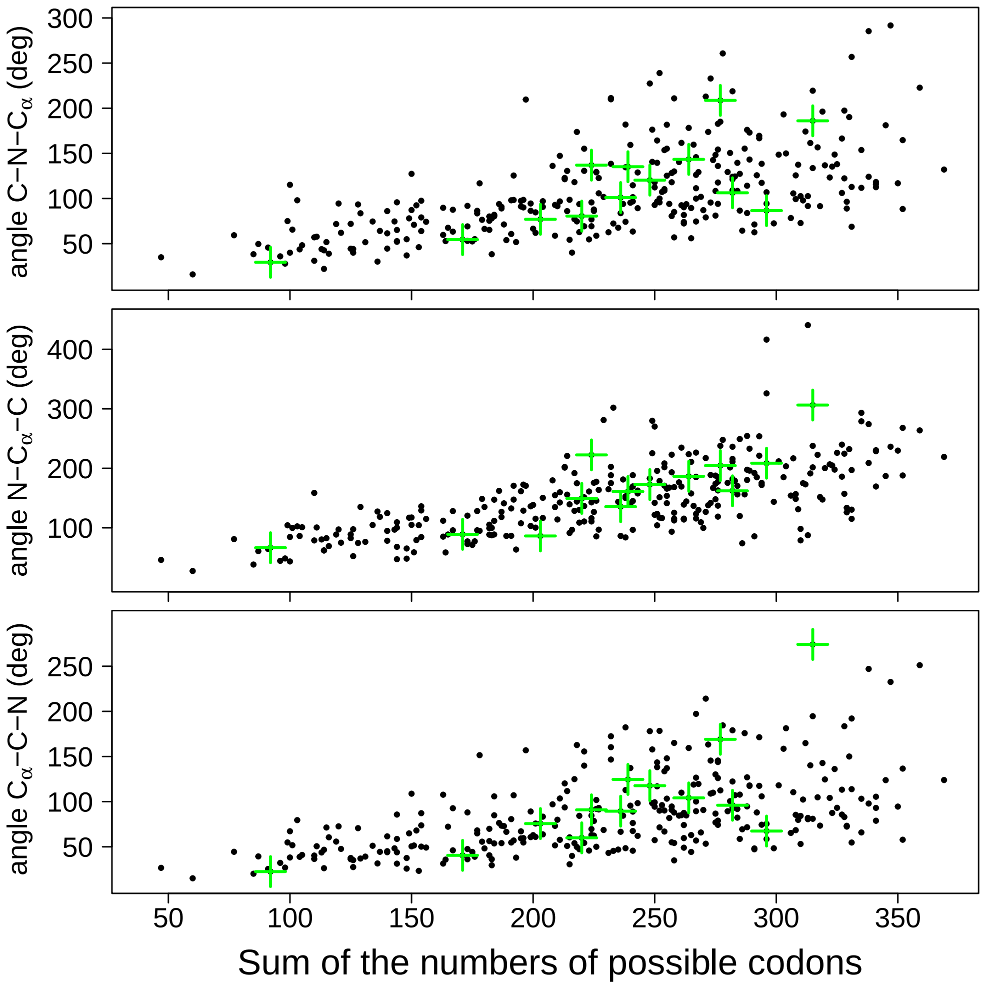


Figure S9: Relationship between the number of possible synonymous codons obtained using the statistics of Table 1 from [6] and the global variation $\Delta\theta$ (Eq. 1 of the main text). Each protein chain in the database is represented by a dot. The 13 protein structures, originating from *E coli* and expressed in *E coli* for the structure determination, are represented by a green cross.

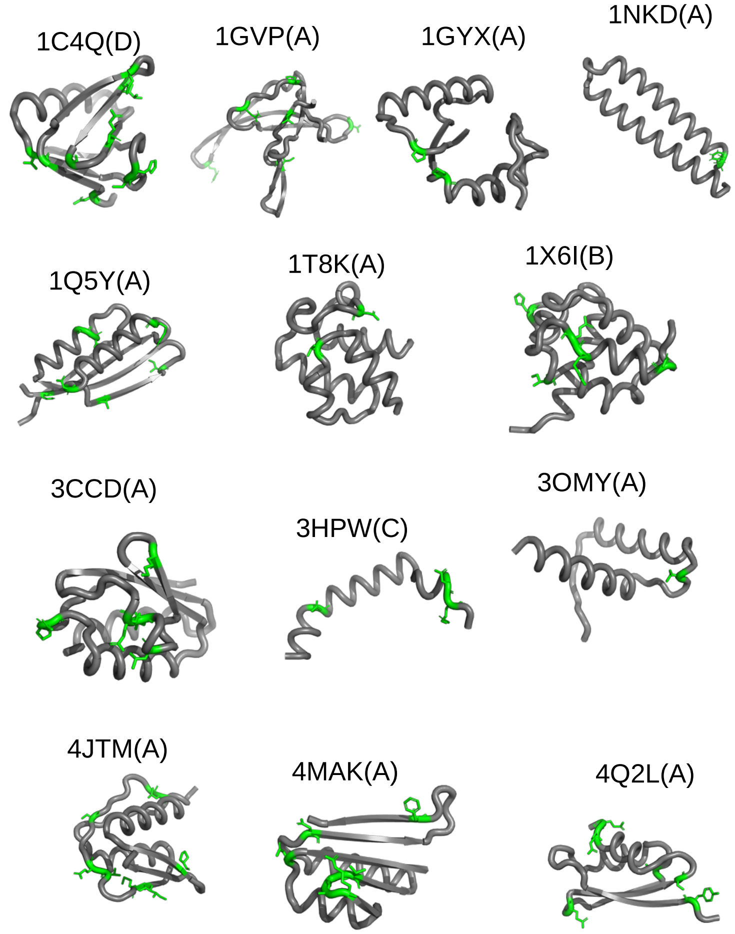


Figure S10: For the 13 protein structures, originating from *E coli* and expressed in *E coli* for the structure determination, the backbone of the selected chain is represented as a tube, labeled by the PDB and chain ids. The residues displaying local variations $|\theta_{i+1} - \theta_i|$ (Eq. 1 of the main text) for bond angles C−N−$C_\alpha$, N−$C_\alpha$−C or C$\alpha$−C−N larger than 6° are drawn in licorice and colored in green.

1C4Q(D) 1C5E(A) 1DP7(P) 1EDM(B) 1EGW(A) 1EZG(A) 1G2R(A) 1G6X(A) 1GUT(A) 1GVP(A) 1GYX(A) 1I27(A) 1I2T(A) 1IQZ(A) 1IRQ(A) 1J2J(B) 1JO0(B) 1JY2(O) 1L9L(A) 1MK0(A) 1NKD(A) 1OAI(A) 1OK0(A) 1PSR(A) 1Q5Y(A) 1Q7L(B) 1R6J(A) 1R7J(A) 1T8K(A) 1U84(A) 1UCR(A) 1UOY(A) 1UTG(A) 1V5I(B) 1V6P(A) 1VBW(A) 1VCC(A) 1W53(A) 1WMH(A) 1X6I(B) 1Y43(A) 1YD0(A) 1YN3(A) 1Z3E(B) 1ZKE(A) 1ZUU(A) 1ZV1(B) 1ZZK(A) 2A26(A) 2AIB(A) 2AYD(A) 2B97(A) 2BAY(A) 2BKF(A) 2CB8(A) 2CC6(A) 2CCQ(A) 2CG7(A) 2CS7(A) 2DS5(A) 2F60(K) 2FCW(B) 2FMA(A) 2G7O(A) 2GOM(A) 2HBA(A) 2HIN(A) 2HS1(A) 2IC6(A) 2IZX(A) 2J5Y(A) 2JKU(A) 2NLS(A) 2O9S(A) 2P5K(A) 2PLX(B) 2QCP(X) 2QFA(B) 2QKV(A) 2V33(A) 2V89(A) 2VC8(A) 2VE8(A) 2WUJ(A) 2WWZ(C) 2XET(A) 2XN6(B) 2XRH(A) 2XTT(A) 2XZ2(A) 2Y5P(D) 2ZW2(A) 3A02(A) 3A0S(A) 3AZD(A) 3B0F(B) 3BQP(A) 3CA7(A) 3CCD(A) 3CIM(A) 3D2Q(A) 3DKM(A) 3DNJ(A) 3DSO(A) 3E7R(L) 3EUN(A) 3FIL(A) 3FJU(B) 3G21(A) 3GE3(C) 3GOE(A) 3H87(C) 3HPW(C) 3IPJ(A) 3LHC(A) 3MAB(A) 3NO7(A) 3ODV(A) 3OMY(A) 3PSM(A) 3RQ9(A) 3SHG(B) 3SJM(A) 3T47(A) 3T7L(A) 3TDU(D) 3TEU(A) 3TVJ(A) 3UFF(A) 3ULJ(A) 3V1A(A) 3VEJ(A) 3VZ9(D) 3W7Y(A) 3W7Y(B) 3WUP(A) 3X34(A) 3ZHI(A) 3ZOQ(B) 3ZR8(X) 3ZZP(A) 4A4J(A) 4A56(A) 4AQO(A) 4CSR(A) 4DRI(B) 4DUQ(A) 4EIC(A) 4EMN(A) 4F87(A) 4FZP(A) 4G3O(A) 4G6T(B) 4GOF(A) 4H4N(A) 4HCS(A) 4HE6(A) 4HGU(A) 4HI8(B) 4HRO(A) 4I0W(A) 4I6R(A) 4IEJ(A) 4J42(A) 4J8C(A) 4JTM(A) 4K12(A) 4K12(B) 4KU0(A) 4L5E(A) 4M8A(A) 4MAK(A) 4NG0(A) 4NPD(A) 4PWW(A) 4Q2L(A) 4QBO(A) 4QXB(B) 4R5R(A) 4RBX(A) 4RGD(A) 4RP3(A) 4RWU(A) 4U5H(A) 4V1G(A) 4WPY(A) 4WZX(A) 4X9Z(A) 4XDX(A) 4XUW(A) 4YNX(A) 4YTW(C) 4ZC3(A) 4ZCE(B) 4ZMK(A) 4ZV5(A) 5AQ0(A) 5AZW(A) 5B1A(U) 5B5I(A) 5COY(A) 5D8V(A) 5EMX(A) 5EQ0(A) 5EU0(B) 5EZU(A) 5GRQ(A) 5H6X(A) 5HUB(A) 5ITM(A) 5JRT(A) 5K2L(A) 5K6D(A) 5KLE(A) 5KXH(B) 5KY0(B) 5KY4(B) 5KY5(B) 5L0V(B) 5L37(C) 5L87(A) 5LEO(A) 5LUS(A) 5M2O(B) 5M97(B) 5MAO(A) 5N41(A) 5NFM(A) 5NW3(A) 5O0U(A) 5O75(A) 5OBT(E) 5OL4(A) 5OXZ(B) 5SVY(A) 5TAB(A) 5TDA(A) 5TW9(A) 5V1V(B) 5V2O(A) 5VGB(B) 5WD9(A) 5X57(A) 5YIU(A) 5YLG(A) 5YP7(A) 5YWR(B) 5ZCY(A) 6ATW(A) 6B29(A) 6BSC(A) 6BSC(B) 6CBU(A) 6CDX(B) 6D0H(B) 6DGA(A) 6E6Q(B) 6ELM(A) 6EXX(A) 6F5Z(C) 6FBQ(A) 6FTO(A) 6FTO(C) 6FU9(A) 6FXA(A) 6G1C(V) 6G6K(A) 6G8Y(A) 6GDJ(A) 6H9U(B) 6HA4(A) 6HBB(A) 6IIP(A) 6IY4(I) 6J4K(B) 6JK2(A) 6KWZ(A) 6L0O(A) 6L0V(E) 6L1P(A) 6MRR(A) 6N9H(A) 6O5K(A) 6OVM(B) 6Q00(A) 6Q6R(G) 6Q9L(A) 6R5J(C) 6RO6(F) 6SCQ(A) 6SID(A) 6SIG(D) 6SQP(B) 6T7O(A) 6T9Q(A) 6TGJ(A) 6TGS(A) 6TIF(B) 6TRJ(A) 6UFE(A) 6UXE(B) 6VG5(B) 6W0V(B) 6XFJ(B) 6XFU(B) 6XIP(C) 6XMI(C) 7CN7(C)

Table S1: List of the 308 X-ray crystallographic structures and protein chains used for the conformation calculations using iBP. The four first characters correspond to the PDB entry and the character in parenthesis corresponds to the chain id.

| Atom name | Definition of atom type | Bond atoms | Bond length (Å) |
|---|---|---|---|
| CH1E | $\alpha$ carbon | C-CH1E | 1.525 |
| C | carbonyl carbon | CH1E-HA | 1.080 |
| O | carbonyl oxygen | CH1E-NH1 | 1.458 |
| OC | C-terminal carbonyl oxygen | CH1E-NH2 | 1.486 |
| HA | H$\alpha$ hydrogen | C-NH1 | 1.329 |
| NH1 | amide nitrogen | C-O | 1.231 |
| NH2 | N-terminal amide | C-OC | 1.249 |
| | nitrogen | H-NH1 | 0.980 |
| H | amide hydrogen | H-NH2 | 0.980 |
| **Bond angle atoms** | **Bond angle value (°)** | **Improper angle atoms** | **Improper angle value (°)** |
| C-CH1E-HA | 108.9914 | C-CH1E-HA-HA | -70.4072 |
| C-CH1E-NH1 | 111.1396 | CH1E-C-NH1-HA | 66.2535 |
| C-CH1E-NH2 | 106.9610 | C-NH1-HA-HA | -70.8745 |
| C-NH1-CH1E | 121.6541 | HA-CH1E-HA-HA | -66.5692 |
| CH1E-C-NH1 | 116.1998 | CH1E-C-NH1-CH1E | 178.0 |
| CH1E-C-O | 120.8258 | O-C-NH1-H | 178.0 |
| CH1E-NH1-H | 119.2367 | C-CH1E-NH1-O | 0.0 |
| C-NH1-H | 119.2489 | NH1-CH1E-C-HA | 119.0 |
| C-NH2-H | 118.1853 | NH2-CH1E-C-HA | 119.0 |
| HA-CH1E-NH1 | 108.0508 | NH1-HA-CH1E-C | 121.0 |
| H-NH2-CH1E | 109.5000 | NH2-HA-CH1E-C | 116.0 |
| H-NH2-H | 107.3000 | C-NH1-CH1E-H | 180.0 |
| NH1-C-O | 122.9907 | CH1E-C-NH1-O | 180.0 |
| NH2-CH1E-HA | 108.4800 | NH2-H-H-CH1E | 41.0 |
| NH2-C-O | 122.6277 | CH1E-OC-C-OC | 178.0 |
| CH1E-C-OC | 118.0611 | | |
| OC-C-OC | 123.3548 | | |

Table S2: Geometric parameters for covalent and improper bonds and angles taken from the force field PARALLHDG (version 5.3) [2].

| Region of Figure S2 | Regions defined in the Ref [7] |
| --- | --- |
| A | $\alpha$ |
| D | $\delta$ |
| d | $\delta$' |
| E | $\epsilon$ |
| P | PII |
| p | PII' |
| Z | $\zeta$ |
| U | unspecified |

Table S3: Correspondence between the regions defined in Figure S2 of supplementary material and the definitions given in the Ref [7]. This correspondence is the same than the one used in Figure S1 of Ref Hollingsworth et al [1].

| AA | N-C$\alpha$-C | C$\alpha$-C-N | C-N-C$\alpha$ | C$\alpha$-C-O | O-C-N | $\omega$ |
|---|---|---|---|---|---|---|
| A | 111.0 ± 2.0 | 116.8 ± 1.3 | 121.3 ± 1.5 | 120.5 ± 1.0 | 122.7 ± 1.0 | 179.1 ± 10.1 |
| R | 111.0 ± 2.2 | 116.8 ± 1.3 | 121.4 ± 1.6 | 120.4 ± 1.0 | 122.7 ± 1.0 | 179.5 ± 9.8 |
| N | 111.4 ± 2.6 | 116.8 ± 1.3 | 121.5 ± 1.7 | 120.4 ± 1.1 | 122.7 ± 1.0 | 179.8 ± 11.6 |
| D | 110.9 ± 2.5 | 116.8 ± 1.3 | 121.5 ± 1.7 | 120.4 ± 1.1 | 122.7 ± 1.0 | 180.1 ± 9.7 |
| C | 110.8 ± 2.3 | 116.7 ± 1.6 | 121.4 ± 1.6 | 120.4 ± 1.1 | 122.9 ± 1.0 | 179.1 ± 11.0 |
| Q | 111.0 ± 2.2 | 116.8 ± 1.3 | 121.3 ± 1.6 | 120.4 ± 1.1 | 122.7 ± 1.0 | 179.2 ± 10.9 |
| E | 111.1 ± 2.1 | 116.9 ± 1.3 | 121.3 ± 1.5 | 120.4 ± 1.0 | 122.7 ± 1.0 | 179.4 ± 10.7 |
| G | 113.2 ± 2.5 | 116.7 ± 1.5 | 121.4 ± 1.5 | 120.5 ± 1.3 | 122.8 ± 1.1 | 180.0 ± 13.2 |
| H | 111.0 ± 2.5 | 116.7 ± 1.4 | 121.5 ± 1.7 | 120.4 ± 1.1 | 122.8 ± 1.0 | 179.4 ± 10.5 |
| I | 109.6 ± 2.3 | 116.7 ± 1.2 | 121.6 ± 1.6 | 120.5 ± 1.0 | 122.7 ± 1.0 | 179.2 ± 8.2 |
| L | 110.8 ± 2.2 | 116.8 ± 1.2 | 121.4 ± 1.6 | 120.4 ± 1.0 | 122.7 ± 1.0 | 178.9 ± 9.4 |
| K | 111.0 ± 2.2 | 116.8 ± 1.3 | 121.5 ± 1.6 | 120.4 ± 1.1 | 122.7 ± 1.1 | 179.4 ± 10.3 |
| M | 110.9 ± 2.2 | 116.8 ± 1.3 | 121.3 ± 1.6 | 120.4 ± 1.1 | 122.7 ± 1.1 | 178.9 ± 9.0 |
| F | 110.7 ± 2.3 | 116.6 ± 1.3 | 121.6 ± 1.7 | 120.5 ± 1.0 | 122.8 ± 1.0 | 178.9 ± 12.5 |
| P | 112.7 ± 2.2 | 116.716 ± 1.6 | 120.1 ± 2.3 | 120.3 ± 1.3 | 122.9 ± 1.1 | 178.5 ± 11.4 |
| S | 111.2 ± 2.3 | 116.7 ± 1.4 | 121.4 ± 1.6 | 120.4 ± 1.1 | 122.8 ± 1.1 | 178.6 ± 11.1 |
| T | 110.7 ± 2.3 | 116.7 ± 1.3 | 121.5 ± 1.6 | 120.5 ± 1.1 | 122.8 ± 1.0 | 178.9 ± 10.7 |
| W | 110.8 ± 2.3 | 116.7 ± 1.4 | 121.5 ± 1.7 | 120.4 ± 1.1 | 122.8 ± 1.1 | 178.9 ± 14.9 |
| Y | 110.8 ± 2.3 | 116.6 ± 1.3 | 121.6 ± 1.7 | 120.5 ± 1.1 | 122.8 ± 1.1 | 178.7 ± 14.5 |
| V | 109.6 ± 2.2 | 116.7 ± 1.2 | 121.7 ± 1.6 | 120.5 ± 1.0 | 122.7 ± 1.0 | 179.0 ± 9.1 |
| Hollingsworth regions | N-C$\alpha$-C | C$\alpha$-C-N | C-N-C$\alpha$ | C$\alpha$-C-O | O-C-N | $\omega$ |
| B | 109.1 ± 2.2 | 116.2 ± 1.2 | 122.2 ± 1.5 | 120.6 ± 1.0 | 123.1 ± 1.0 | 178.0 ± 14.0 |
| P | 110.6 ± 2.0 | 116.2 ± 1.3 | 121.0 ± 1.7 | 120.9 ± 1.1 | 123.0 ± 1.1 | 177.8 ± 15.9 |
| p | 112.0 ± 2.4 | 116.0 ± 1.6 | 121.4 ± 1.7 | 121.1 ± 1.3 | 122.8 ± 1.1 | 181.7 ± 21.1 |
| E | 111.2 ± 2.5 | 116.1 ± 1.5 | 121.8 ± 2.1 | 121.0 ± 1.2 | 123.0 ± 1.1 | 181.1 ± 17.5 |
| U | 110.1 ± 3.4 | 117.3 ± 3.2 | 121.8 ± 1.9 | 120.0 ± 1.4 | 122.2 ± 2.9 | 181.3 ± 19.4 |
| Z | 108.9 ± 2.5 | 116.6 ± 1.3 | 122.4 ± 1.5 | 121.0 ± 1.0 | 122.3 ± 1.1 | 183.7 ± 16.2 |
| d | 113.9 ± 2.2 | 117.1 ± 1.4 | 122.3 ± 1.6 | 120.2 ± 1.3 | 122.6 ± 1.0 | 179.4 ± 6.4 |
| g | 109.9 ± 2.6 | 115.4 ± 1.2 | 121.8 ± 1.4 | 121.6 ± 1.1 | 122.9 ± 1.0 | 183.3 ± 7.5 |
| D | 113.2 ± 1.9 | 117.5 ± 1.2 | 121.7 ± 1.6 | 119.8 ± 1.1 | 122.6 ± 1.0 | 180.9 ± 6.7 |
| A | 111.6 ± 1.7 | 117.1 ± 1.1 | 120.7 ± 1.4 | 120.3 ± 1.0 | 122.5 ± 1.0 | 179.7 ± 4.2 |
| G | 113.7 ± 2.9 | 117.1 ± 1.4 | 124.3 ± 2.6 | 120.0 ± 1.2 | 122.8 ± 1.3 | 181.1 ± 7.0 |

Table S4: Average and standard deviation values for the bond angles between backbone heavy atoms and torsion angle $\omega$. These values corresponds to the ones used in the plot of Figure 2 in the main text and were calculated according to the amino acid type of the atom C$\alpha$ of the analyzed residue, and according to the Hollingsworth region defined from the $\phi$ and $\psi$ torsion angles of the same residue.
